# Multi-species analysis of inflammatory response elements reveals ancient and lineage-specific contributions of transposable elements to NF-κB binding

**DOI:** 10.1101/2022.10.25.513724

**Authors:** Liangxi Wang, Tiegh Taylor, Kumaragurubaran Rathnakumar, Nadiya Khyzha, Minggao Liang, Azad Alizada, Laura F Campitelli, Sara E Pour, Zain M Patel, Lina Antounians, Ian C Tobias, Timothy Hughes, Sushmita Roy, Jennifer A Mitchell, Jason E Fish, Michael D Wilson

## Abstract

Transposable elements (TEs) provide a source of transcription factor binding sites that can rewire conserved gene regulatory networks. NF-κB is an evolutionary conserved transcription factor complex primarily involved in innate immunity and inflammation. The extent to which TEs have contributed to NF-κB responses during mammalian evolution is not well established. Here we performed a multi-species analysis of TEs bound by the NF-κB subunit RELA (also known as p65) in response to the proinflammatory cytokine TNF. By comparing RELA ChIP-seq data from TNF-stimulated primary aortic endothelial cells isolated from human, mouse and cow, we found that 55 TE subfamilies were associated with RELA bound regions. These RELA-bound transposons possess active epigenetic features and reside near TNF-responsive genes. A prominent example of lineage-specific contribution of transposons comes from the bovine SINE subfamilies Bov-tA1/2/3 which collectively contributed over 14,000 RELA bound regions in cow. By comparing RELA binding data across species, we also found several examples of RELA motif-bearing TEs that colonized the genome prior to the divergence of the three species and contributed to species-specific RELA binding. For example, we found human RELA bound MER81 instances were enriched for the interferon gamma pathway and demonstrated that one RELA bound MER81 element can control the TNF-induced expression of Interferon Gamma Receptor 2 (*IFNGR2*). Using ancestral reconstructions, we found that RELA containing MER81 instances rapidly decayed during early primate evolution (> 50 million years ago (MYA)) before stabilizing since the separation of Old World monkeys (< 50 MYA). Taken together, our results suggest ancient and lineage-specific transposon subfamilies contributed to mammalian NF-κB regulatory networks.

## INTRODUCTION

Transposable elements (TEs) are known to play a crucial role in shaping gene regulation by serving as source of novel *cis*-regulatory elements (CREs) that can affect the expression of nearby genes (Chuong et al. 2017; Pontis et al. 2019; Sundaram and Wysocka 2020; Hermant and Torres-Padilla 2021; Senft and Macfarlan 2021; Fueyo et al. 2022). One common process by which TEs would be exapted as CREs involves the acquisition of regulatory function through mutations that create or modify transcription factor (TF) binding sites within the TE sequence.

Identifying which TFs interact with TEs and under what conditions is an ongoing challenge that requires a range of experimental and computational approaches (Bourque et al. 2008; Macfarlan et al. 2011; Emera and Wagner 2012; Jacques et al. 2013; Ward et al. 2013; Sundaram et al. 2014; Ito et al. 2017; Fuentes et al. 2018). Chromatin immunoprecipitation followed by DNA sequencing (ChIP-seq) of an ever growing number of TFs in different cell types, species, and conditions are revealing repeat-associated binding sites whereby reproducible TF-DNA interactions fall within subtypes of major repeat families (Bourque et al. 2008).

Coincident with their discovery, repeat associated binding sites (RABS) have been shown to contribute to gene regulatory networks underlying a myriad of processes including early developmental gene regulation, pregnancy, circadian rhythm and innate immunity (Kunarso et al. 2010; Xie et al. 2010; Lynch et al. 2011; Notwell et al. 2015; Chuong et al. 2016; Trizzino et al. 2017; Fuentes et al. 2018; Pontis et al. 2019; Todd et al. 2019; Bogdan et al. 2020; Judd et al. 2021; Kelly et al. 2022). Further insights into the regulatory potential of TEs are also coming from high-throughput experiments that perturb the activity of young TE families using *CRISPR-Cas9* (e.g. (Fuentes et al. 2018; Pontis et al. 2019)), as well as massively parallel reporter assays (e.g. (Du et al. 2022)).

Nuclear factor kappa-light-chain-enhancer of activated B cells (NF-κB) is an essential mediator of innate immune/inflammatory responses (Zhang et al. 2017), whose activity has been conserved across metazoans (Sen and Baltimore 1986; Satriano and Schlondorff 1994; Correa et al. 2004; Hetru and Hoffmann 2009). In vertebrates, NF-κB is composed of hetero-dimers of five proteins: NFKB1 (also known as p50), NFKB2 (also known as p52), RELA (also known as p65), RELB and REL, with the most predominant combinations being NFKB1:RELA which can be found in most cell types (Bhatt and Ghosh 2014). The NFKB1:RELA complex mediates canonical NF-κB signaling pathways. This process relies on NFKB1:RELA being sequestered in its inactive form in the cytoplasm through its association with the inhibitory IκB proteins under unstimulated conditions. In response to upstream stimuli, such as the pro-inflammatory cytokine tumor necrosis factor (TNF), the IκB inhibitors are degraded and NF-κB rapidly translocates into the nucleus where it binds promoter-proximal and promoter-distal CREs that drive the expression of multiple innate immunity/inflammation related genes (Zhang et al. 2017).

While the core protein components of NF-κB are often shared across cell types, the binding of NF-κB to the genome is constrained by cell-type specific chromatin states (Natoli et al. 2005; Ostuni et al. 2013). For example, abundant NF-κB binding occurs at the pre-existing SPI1 (PU.1) transcription factor bound regions in macrophages (Kaikkonen et al. 2013) whereas NF-κB binding in aortic endothelial cells occurs at ERG and JUN bound regions (Hogan et al. 2017). In addition to comparisons of NF-κB binding across cell types, comparing its genome-wide binding across species can reveal conserved NF-κB regulatory principles and highlight functional CREs that drive NF-κB target gene expression (Alizada et al. 2021). For example, an inter-species comparison of NF-κB binding in primary aortic endothelial cells stimulated with TNF for 45 minutes found that conserved orthologous NF-κB bound regions were found to be enriched near NF-κB target genes, coincided with increased RELA binding and chromatin accessibility, increased post-translational histone modifications associated with active promoters and enhancers (H3K27ac and H3K4me2), increased RNA polymerase II activity in chromatin run-on assays (ChRO-seq) (Alizada et al. 2021). Furthermore, this comparative epigenomic analysis of NF-κB showed that conserved orthologous NF-κB bound regions are active across diverse cell types and overlap non-coding genetic variation associated with both inflammatory and cardiovascular phenotypes.

Pioneering work using circular chromosome conformation capture (4C) coupled with RELA ChIP in HeLa cells cloned and validated three sequences involved in interchromosomal interactions of the interferon beta (*IFNB*) promoter. All three of these cloned regions contained *Alu* repeat sequences containing DNA motifs similar to the NF-κB consensus motif (Apostolou and Thanos 2008). Subsequent cloning of 366 sequences obtain by RELA ChIP in HeLa cells combined with the analysis of ENCODE RELA ChIP-seq (26-bp single end data) from the lymphoblast cell line GM12878 led to the conclusion that ∼11% of NF-κB bound regions contain an *Alu* derived NF-κB motif (Antonaki et al. 2011). However, the extent to which specific subfamilies of repetitive elements contribute to NF-κB binding in primary human cells and cells from other mammalian species remains to be characterized. To address the contribution of TEs to NF-κB binding during mammalian evolution we performed a comparative genomic analysis of NF-κB binding induced by TNF stimulation of primary human, mouse and cow aortic endothelial cells.

## RESULTS

### Transposable elements contribute to mammalian NF-κB bound regions

To identify the contribution of transposable elements to NF-κB bound regions relative to other transcription factors, we first interrogated genome-wide binding data for 435 human and 266 mouse TFs found in the Gene Transcription Regulation Database (GTRD) (Kolmykov et al. 2021). The GTRD database contains a wide variety of uniformly processed TF ChIP-seq data across many contexts and cell types, including 47 datasets from 10 cell types for the NF-κB subunit RELA (Kolmykov et al. 2021). We calculated the fraction of RABS by overlapping ChIP-seq peak summits with RepeatMasker annotated TEs (Bao et al. 2015). While transcription factor bound regions are globally depleted for TEs (median TE overlap was 25% for human and 18% for mouse), TE-derived RELA bound regions were above the median of all transcription factors (84th percentile and 77th percentile of all investigated TFs for human and mouse, respectively), with a median overlap of 33% and 21% for human and mouse respectively (Supplementary Figure 1A,B). This result suggests that TEs contributed substantially to the dissemination of human and mouse NF-κB bound regions.

To study the evolution of TE-derived RELA binding, we then focused our analysis on our previously published work that mapped RELA binding by ChIP-seq in human, mouse and cow (Alizada et al. 2021). This comparative epigenomics study was performed in primary human, cow and mouse aortic endothelial cells (HAEC, BAEC, and MAEC respectively) treated acutely (45 min) with a species-matched pro-inflammatory cytokine tumor necrosis factor (TNF). This study also profiled a karyotypically normal immortalized cell line teloHAEC that shows a similar NF-κB response to TNF as HAECs and is amenable to *CRISPR-Cas9* genome editing (Alizada et al. 2021) (Figure 1). To facilitate direct comparisons between this study and the previous study, we aligned RELA ChIP-seq data to human (hg19), mouse (mm10) and bovine (bosTau6) genome builds found in the multiple genome alignment (Ensembl 70 release) used by Alizada et al. 2021. This dataset gave a significant number of RELA bound regions for each species after 45 minutes of TNF treatment (identified by using MACS2 (Zhang et al. 2008) to call peaks on the treated sample using the untreated sample as input; human: 59,735; mouse: 23,623; and cow: 93,511). Note that there were only 383 RELA peaks unique to hg19 and 420 peaks unique to the hg38 genome build suggesting the choice of genome build does not significantly affect our results. While our use of 100bp single end reads will be insufficient for mapping the youngest transposon families due to their highly repetitive nature (Sexton and Han 2019), we reason that this unique comparative epigenomic dataset, which includes ATAC-seq (all three species) and H3K4me2 and H3K27ac (human), is well-suited for comparing mammalian repeat associated NF-κB binding between species..

**Figure 1.**
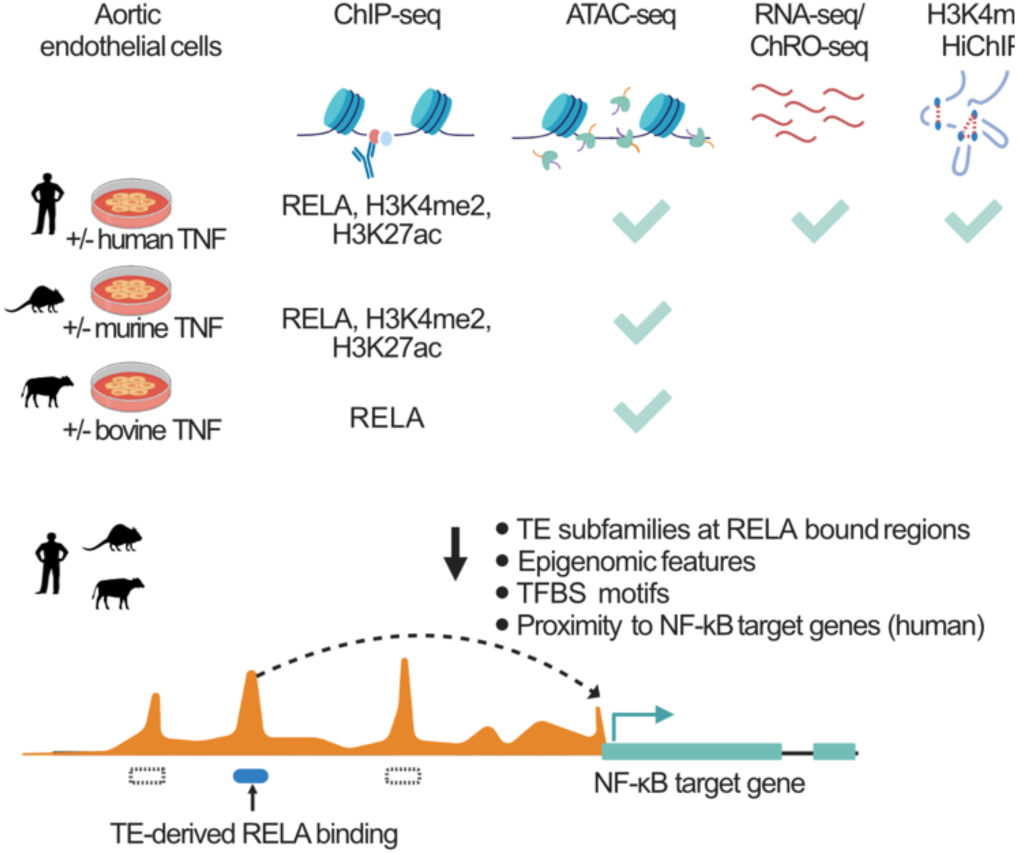
Schematic of the experimental data and computational workflow. Primary aortic endothelial cells from three mammalian species were cultured with and without 45-minute treatment of the pro-inflammatory cytokine TNF. These cells were subjected to chromatin immunoprecipitation followed by DNA sequencing (ChIP-seq) assays using antibodies against RELA, H3K4me2 and H3K27ac. Chromatin accessibility was assessed using ATAC-seq. Immortalized human aortic endothelial cells (TeloHAEC) were used for ChRO-seq and H3K4me3 HiChIP. Here we performed H3K4me3 HiChIP to identify active enhancer-promoter connections in human. Transposable element derived RELA-bound regions were computationally determined. In particular, the contribution of transposable elements was examined across three species and four cell types. These elements exhibit unique sequence and epigenetic features and functionally associate with target gene expression. This figure was created with BioRender.com.

Compared to the median TE overlap of RELA in the GTRD database, we observed a slightly lower overlapping fraction of TEs within RELA-bound regions in human and mouse aortic endothelial cells (19% and 12% respectively). In contrast, we observed a striking fraction (∼31%) of RELA bound regions in cow to be TE derived (Supplementary Figure 1C). To investigate this further we compared the proportions of the four major TE classes (LINE, SINE, DNA transposon, LTR) that overlap RELA peak summits to the background expectation defined by the genome coverage of each class. For all species the observed TE class proportions were significantly different from background expectations (chi-squared test, corrected p values < 1×10^-7^ for all tests; Supplementary Figure 1C). The most striking example of this difference was for the SINEs in cow, which overlapped 20,188 RELA peaks, at a frequency of 22% (genome background of 17%; chi-squared test, 1.3-fold enriched, corrected p value= 2.6×10^-267^). Restricting the analysis to TE-derived sequences, we observed that the LTR class was close to two-fold enriched within RELA-bound regions for both human and mouse, but not in cow (chi-squared test; human: 1.8-fold enriched, p value< 2×10^-308^; mouse: 1.8-fold enriched, p value< 2×10^-308^; cow: 0.99-fold enriched, p value=0.53. Supplementary Figure 1D).

We compared TE derived RELA bound regions from each species using whole genome multiple sequence alignments which allowed us to identify orthologous and species specific NF-κB binding events (Alizada et al. 2021). As expected, we found most TE-derived RELA-bound regions to be species-specific (chi-squared test; human: 90%, mouse: 92%, cow: 95%; corrected p values <3.6×10^-207^ for all three species; Supplementary Figure 2A).

We then took a closer look into the instances where TE-derived RELA bound regions occurred in the context of conserved orthologous RELA binding. Interestingly, the majority of conserved orthologous TE-derived RELA-bound regions were classified as being TE-derived in only one species (human: 810/1026; mouse:127/255 and cow: 920/1410). Two explanations for this observation are that: 1) there is an orthologous TE that was not annotated by RepeatMasker in the second species; or 2) a conserved orthologous RELA binding site turned over into an adjacent TE-derived DNA sequence. To explore these possibilities, we split our human conserved RELA binding events into three categories: Category 1 – conserved orthologous RELA binding is not related to TEs (n=16,522); Category 2 – conserved orthologous RELA binding involved TEs in one or more species (n=221); and Category 3 – conserved orthologous RELA binding was TE derived only in human (n=899; Supplemental Figure S2B).

Both Category 1 and Category 2 regions showed high DNA constraint. In contrast, Category 3 showed lower DNA constraint that was comparable to the DNA constraint observed for human-specific RELA binding. To investigate this further, we extracted and realigned all human RELA-bound sequences (+/-500 bp from the summit) that had conserved orthologous RELA binding in at least one other species. We then compared the base pair distance between human RELA peak summits and the ‘lifted-over’ summit position from mouse or cow.

Consistent with an enrichment of micro-turnover events in Category 3, we found that the lifted over RELA peak summits in Category 3 were further apart than what was observed for Category 1 or 2 (Supplemental Figure 2C). We then measured the base pair distance of lifted-over mouse or cow RELA peak summits to the nearest human TE. As would be expected for conserved orthologous TE-derived RELA binding events, we observed that ∼75% of the mouse/cow RELA peak summits in Category 2 were within 25bp of a human TE. In contrast, ∼50% of the RELA peak summits in Category 3 were within 25bp of a human TE (Supplemental Figure 2D). This indicates that Category 3 events likely represent a combination of missing TE annotations in addition to micro-turnovers into adjacent TEs, several of which are primate lineage-specific TEs (Supplemental Figure 2E, F).

To further investigate the TE features that contributed to TE-derived RELA-bound regions, we annotated each RELA peak summit with its corresponding TE subfamily using RepeatMasker (Figure 2A). We identified 55 significantly enriched TE subfamilies among all three species, 7 of which were shared in at least one additional species (Figure 2A,B). These enriched subfamilies collectively make up 9% (1020/11365), 13% (369/2903) and 52% (15475/29576) of TE-derived RELA-bound regions in human, mouse and cow, respectively. Notably, 22 subfamilies contained canonical RELA motifs in their consensus sequences (Figure 2B). LTRs account for the majority of TE subfamilies contributing RELA-bound regions in the human and mouse experiments (human: 68%, mouse: 65%, and cow: 2%). The only LTR that was enriched for RELA binding in all three species was MER21C. A notable example of a primate LTR subfamily associated with RELA binding is MER41B, a TE subfamily previously shown by Chuong *et al*. 2016 to be bound by interferon gamma (IFNG)-inducible TFs STAT1 and IRF1 in immortalized human cell lines and primary macrophages. In contrast to human and mouse, the SINE class of TEs makes up the majority (68%) of TE-derived RELA-bound regions in the cow genome. Four SINE subfamilies Bov-tA1, Bov-tA2, Bov-tA3, and SINE2-1_BT account for 14,914 (74%) of these RELA-bound SINEs. The presence of a RELA motif in the consensus sequence of Bov-tA3, which overlaps 2,252 RELA binding events, provides a plausible explanation for these SINE-derived RELA-bound regions (Figure 2B).

**Figure 2.**
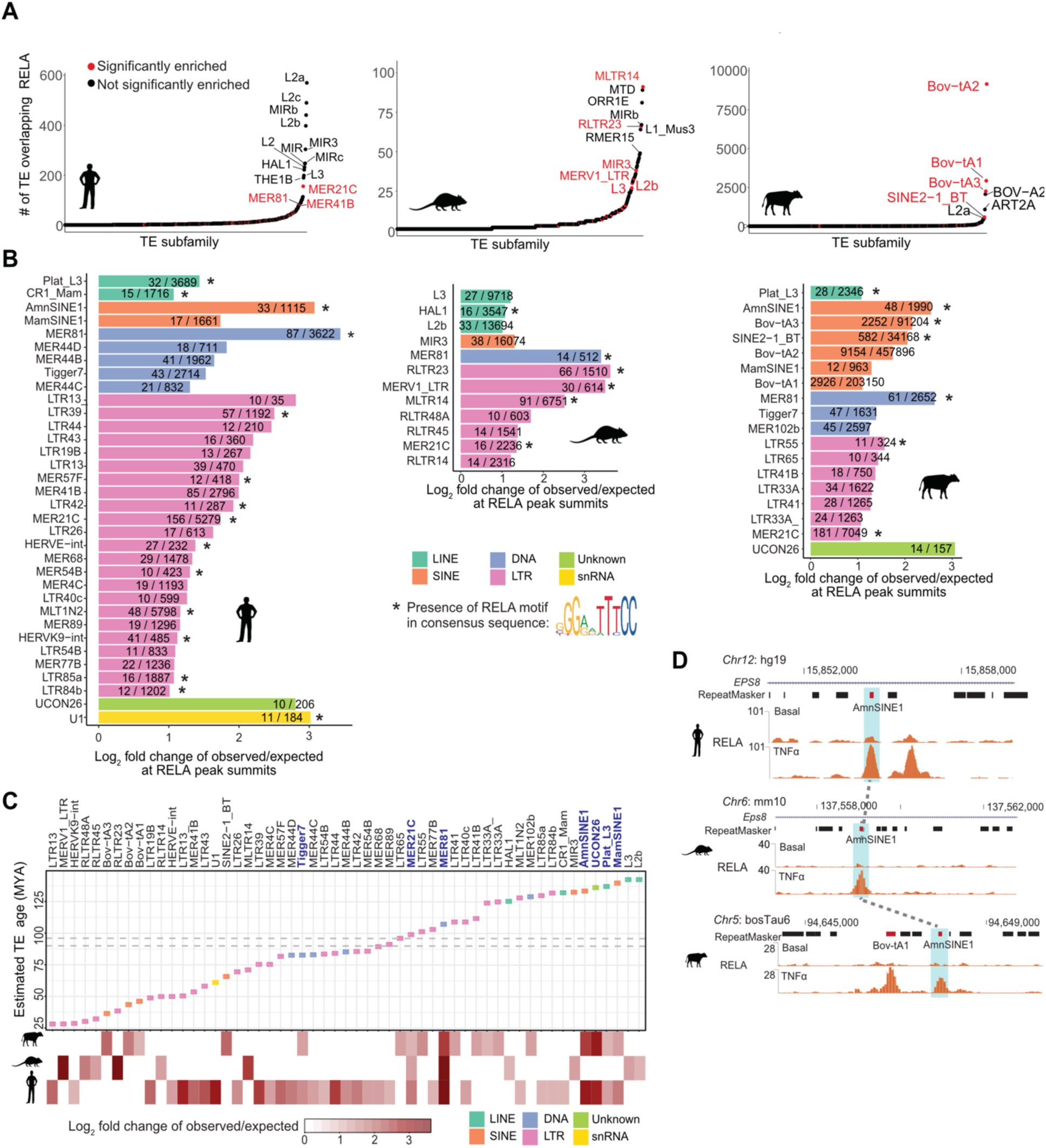
Transposable elements contribute to mammalian NF-κB bound regions. RELA ChIP-seq data with/without TNF treatment for 45 min from human/mouse/cow aortic endothelial cells were used to characterize RELA-bound regions in the three species. A) Ranked overlap values for each TE subfamily with RELA peak summits from the three mammals. Significantly enriched TE subfamilies found in panel B were colored in red. B) TE subfamilies that were significantly enriched within RELA-bound regions in human, mouse and cow. The enrichment values were calculated by comparing observed and expected/simulated overlap events at RELA peak summits. The cutoffs to define enriched TE subfamilies: FDR < 0.05, log_2_ fold change > 1 and the number of observed overlap events >= 10. Observed overlap number/ total number of each TE subfamily is displayed for each bar. Bars marked with an asterisk indicate the presence of canonical RELA motifs in the consensus sequence. C) Estimated evolutionary age of all the enriched TE subfamilies. The top panel depicts the median of estimated age of these elements, while the bottom panel presents enrichment values. Dashed gray lines represent estimated human-mouse and mouse-cow divergence time obtained from TIMETREE database (Kumar et al. 2017). Bars marked with an asterisk indicate the presence of canonical RELA motifs in the consensus sequence. TE subfamilies shared by more than one species are highlighted in blue. D) An example (AmnSINE1) of a three-species-conserved TE-derived RELA-bound region. Red block from RepeatMasker track represents RELA-bound TE and light blue shades show orthologous regions of the three species.

To broadly classify TEs into ancient and lineage specific categories we approximated the evolutionary age of the 55 enriched TE subfamilies using the sequence divergence of subfamily instances relative to their consensus sequence. Seven of the enriched TE subfamilies were considered ancient (Tigger7, MER21C, MER81, AmnSINE1, UCON26, Plat_L3, MamSINE1) and likely to have colonized the mammalian genome near or prior to the divergence of the three species (∼90 MYA (Kumar et al. 2017)) Figure 2C). Of these ancient sub-families, AmnSINE1, MER81, and UCON26 were enriched for NF-κB binding in at least two of the three species. For example, AmnSINE1 colonized the mammalian genome before the radiation of amniote animals (Nishihara et al. 2006) and has been shown to contribute CREs to conserved gene regulatory networks active during brain development (Sasaki et al. 2008; Hirakawa et al. 2009). AmnSINE1 elements were significantly enriched for human and cow RELA binding. Although this TE was not considered to be significantly enriched in mouse, we found 9 RELA bound instances (4.6-fold enrichment; one example is shown in Figure 2D). Another striking example of an ancient RELA associated TE was MER81, a 114 bp non-autonomous DNA transposon derived from MER91 sequence (Bao et al. 2015). MER81 was enriched for RELA binding in all three species (10.9-fold change in human, 10.6-fold change in mouse and 6.2-fold change in cow). Thus both ancient and lineage specific TE subfamilies have contributed to the NF-κB binding repertoire.

### RELA-bound TEs possess active enhancer features for target gene regulation

To explore the regulatory potential of the TE-derived RELA-bound regions, we assessed epigenomic proxies for active CREs in TNF-treated and untreated endothelial cells in each species. Starting with the 55 TE subfamilies in human, we investigated the dynamic changes in RELA binding, H3K27ac (active enhancers/promoters), H3K4me2 (active or latent enhancers/promoters), and chromatin accessibility (ATAC-seq) before and 45 minutes after TNF treatment (Alizada et al. 2021). We first subcategorized these elements into “old-TEs” and “young-TEs’’ where “old” TE subfamilies were designated based on their estimated colonization of mammalian genomes being prior to the divergence of the selected mammals (90 MYA). As conserved orthologous RELA binding is closely linked to high RELA binding, H3K27ac signal, enhancer activity and proximity to NF-κB target genes (Alizada et al. 2021), we included conserved RELA-bound regions in this analysis (regardless of whether or not they were derived from TEs) as a “positive control” reference set. The remaining human-specific bound regions (regardless of whether or not they were derived from TEs) were included as a benchmark for baseline enhancer features at RELA bound regions.

We found that the old and young TE-derived RELA-bound regions were associated with high RELA binding signals after TNF treatment and were comparable to that of conserved RELA bound regions (Figure 3A; Supplementary Figure 3). Increased chromatin accessibility was also observed for both TE-derived groups post TNF treatment. While a clear signature of active CREs, characterized by H3K4me2 and H3K27ac, was associated with TE-derived groups, this TE-derived epigenomic signal was far lower than what was observed for the conserved RELA bound regions and slightly lower than the human-specific group. Importantly, evidence for TE-derived RELA bound regions being associated with active CRE characteristics was also observed for mouse and cow TE-derived RELA bound regions (Supplementary Figure 4).

**Figure 3.**
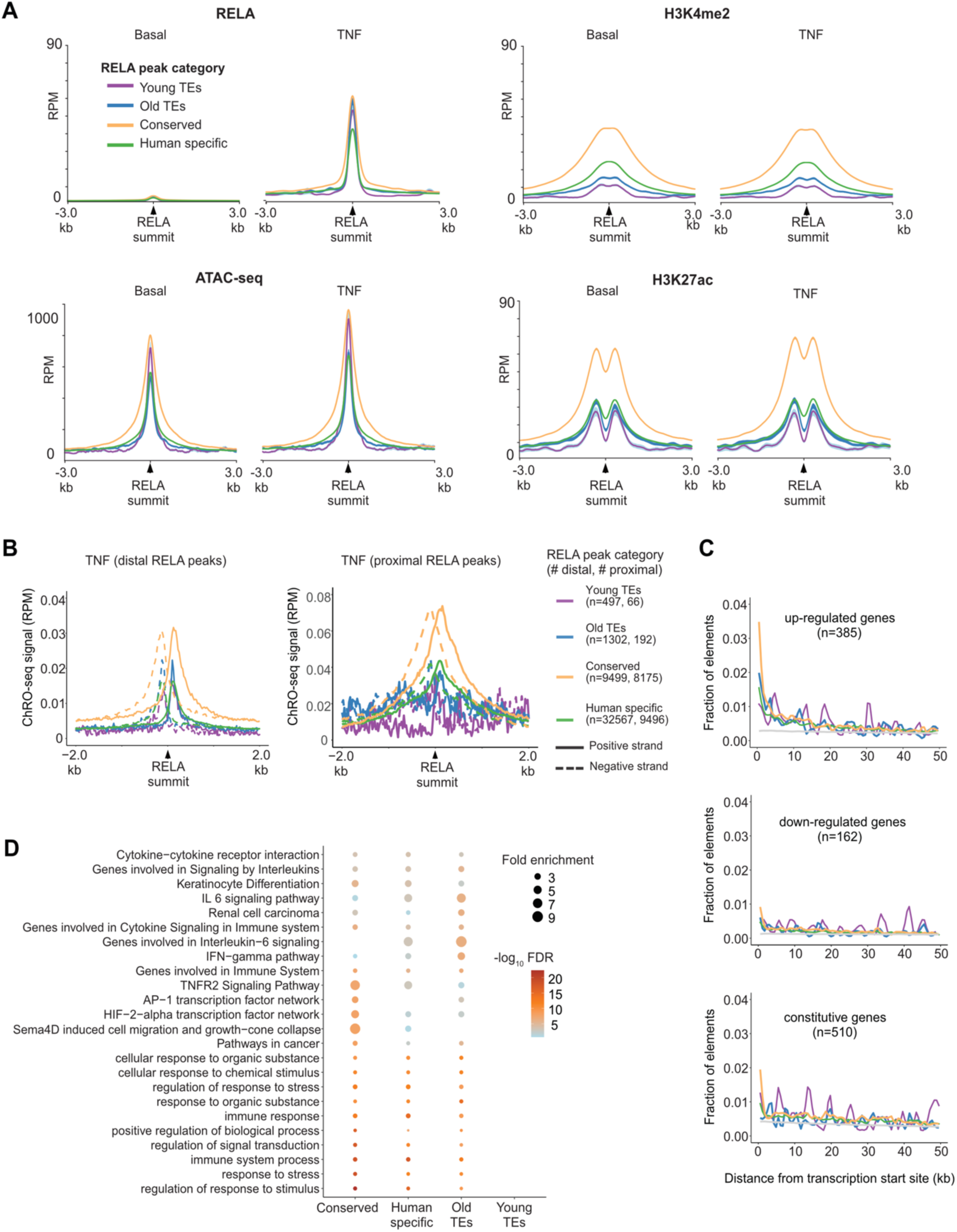
RELA-bound TEs possess active enhancer features for target gene regulation. Four categories of RELA-bound regions were interrogated: young TE-derived RELA-bound regions, old TE-derived RELA-bound regions, conserved RELA-bound regions and human-specific RELA-bound regions. A) Epigenetic profile around RELA-bound TEs before and after TNF stimulation. Included are aggregated signals of RELA ChIP-seq, H3K4me2 and H3K27ac ChIP-seq and ATAC-seq data from human aortic endothelial cells. B) eRNA signal from ChRO-seq of TeloHAEC around RELA-bound TEs that are distal or proximal to gene TSS with TNF stimulation for 45 min. The cutoff for distal/ proximal elements: 3 kb relative to gene TSS. C) The distribution of RELA-bound TEs near up-regulated, down-regulated and constitutive genes. The three groups of genes were defined using matched RNA-seq data (see Methods). “Background” represents the distribution of all the genomic TEs near the three gene groups, shown in gray. D) Functional enrichment of RELA-bound TEs by GREAT (McLean et al. 2010).

To further assess the enhancer features of TE-derived subsets in human, we analyzed data from a genome-wide global RNA polymerase run-on assay, which we previously generated from a karyotypically normal, immortalized human aortic endothelial cell line (TeloHAEC) (Fish et al. 2017; Alizada et al. 2021). This profiling was achieved using ChRO-seq (Chromatin Run-On sequencing (Chu et al. 2018)) which reveals bi-directional transcription of enhancer RNAs (eRNAs), a hallmark of active enhancers (Kaikkonen et al. 2013). Conserved RELA-bound regions showed the highest eRNA signal followed by old TE-derived RELA-bound regions. The majority of the two TE categories reside within the distal genomic regions (more than 3 kb away from gene TSS). We found a clear pattern of bi-directional transcription of eRNAs in all four RELA groups at both distal (> 3 kb) and proximal (<= 3 kb) regions relative to the gene TSS (merely one TSS was considered per gene; Figure 3B). Overall, this analysis supports the possibility that transposons contribute active enhancers which are involved in NF-κB mediated gene regulation.

To have a functional impact, we would expect TE-derived RELA bound regions to be found in the vicinity of NF-κB target genes. To address this question we reanalyzed the RNA-seq data that was paired with the RELA ChIP-seq data obtained from primary human aortic endothelial cells before and after TNF treatment (Alizada et al. 2021). We assessed the likelihood that RELA-bound TEs influenced NF-κB target gene expression by comparing the distribution of the four categories of RELA bound regions around up-regulated, down-regulated, and constitutive genes (Figure 3C; see Methods). We observed a significant enrichment of RELA-bound regions near the promoters of up-regulated genes compared to constitutive genes, with the old TE-derived RELA group showing the highest enrichments, followed by the conserved RELA group (binomial test for occurrences within 10kb relative to up-regulated vs constitutive gene TSS. Old TEs: 3.6-fold, 6.9×10^-9^; Conserved: 1.8-fold, p-value< 2.2×10^-16^; Human specific: 1.6-fold, p-value< 2.2×10^-16^; and Young TEs: 0.9-fold, p-value=0.85).

Accordingly, TE-derived RELA-bound regions were enriched for annotated immune/inflammatory functions (Figure 3D). While there was no enrichment of the young TE-derived RELA group (n=561), we found comparable enrichment signals in functionally related terms for the other three groups. We noted several terms particularly enriched in the old TE-derived RELA group, including “IFN-gamma pathway” (binomial FDR of 2.2×10^-7^) and “IL6 signaling pathway” (binomial FDR of 1.06×10^-6^). These observations suggest the possibility that ancient TE-derived RELA bound elements have preferentially contributed to IFN-gamma/IL6 signaling during human evolution.

While linear proximity between a *cis*-regulatory element and a promoter is associated with potential function, three-dimensional chromatin interactions provide additional direct evidence of a physical connection between distal CREs and gene promoters (Chepelev et al. 2012; Mifsud et al. 2015; Fulco et al. 2019). To profile the interaction of TE-derived RELA bound regions with gene promoters, we performed HiChIP (Fang et al. 2016; Mumbach et al. 2016) in TeloHAECs before and 45 min after TNF stimulation using an antibody against H3K4me3, a post-translational histone modification enriched at active gene promoters. We found a union of 19,372 significant chromatin interactions before and 45 min after TNF stimulation, 1605 of which involved NF-κB up-regulated genes. Considering the portion of distal elements (at least 5 kb away from any gene TSS) from the four categories, conserved RELA-bound regions have the highest fraction of overlap with the distal anchor from the identified loop sets (28%), followed by old TE-derived (15%), human-specific (15%) and young TE-derived (11%) RELA bound regions. We next focused on the subset of chromatin interactions (n=1605) that connect 1180 distal RELA bound regions and 224 NF-κB up-regulated genes. A subset of these elements (186 out of 1180) were derived from TEs, 39 of which belong to the significantly enriched transposons identified in Figure 2B (Supplementary Material 1). Overall, these 3D proximity results between transposons and genes further support the possibility that TE-derived RELA-bound regions have been incorporated into NF-κB gene regulatory networks.

### TE-derived RELA bound regions coincide with cell-type specific TF motifs

The NF-κB signaling pathway is active across diverse cell types and involves common and cell-type specific target gene expression (Tabruyn and Griffioen 2007; Gerondakis and Siebenlist 2010; Liu et al. 2017). While cell-type specific NF-κB binding is known to occur in collaboration with cell-type specific TFs (Kaikkonen et al. 2013; Hogan et al. 2017; Alizada et al. 2021), it was not known if specific cell types would be enriched for distinct TE subfamilies at NF-kB bound regions. To address this question, we reanalyzed data from three additional cell types which all showed an appreciable number of RELA bound regions after TNF stimulation: human umbilical vein endothelial cells (HUVECs; n = 12,442 (Brown et al. 2014)), lymphoblasts (LCLs; n = 52563 (Kasowski et al. 2010)) and adipocytes (n = 58,043 (Schmidt et al. 2015)) (Figure 4A). After calculating the overlap of RELA peak summits with RepeatMasker annotated TE subfamilies, we observed both cell-specific and cell-shared enrichments (Figure 4B and 4C). We found that HUVECs showed a subset of TE-derived RELA binding that we observed in our human aortic endothelial cell (HAEC) data. In contrast we found enrichments for multiple TE subfamilies unique to adipocytes, as well as a few highly enriched endogenous retroviruses specific to LCLs (e.g., LTR43, LTR43B, MER57E1, and MER73). Notably MER81 and AmnSINE1 subfamilies, which we previously identified as being enriched at RELA bound regions in human, mouse and cow aortic endothelial cells, were also found to be enriched across multiple cell types. MER81 showed the highest enrichment values in almost all four selected cell types (10.6-, 9.2-, 18.4-, and 8.6-fold enrichment for HAEC, HUVEC, Adipocytes, and LCL, respectively). AmnSINE1 shows both high conservation degree and cross-cell usage of RELA-bound instances when compared across cell type and species (with 60% and 53% respectively; Supplementary Figure 5).

**Figure 4.**
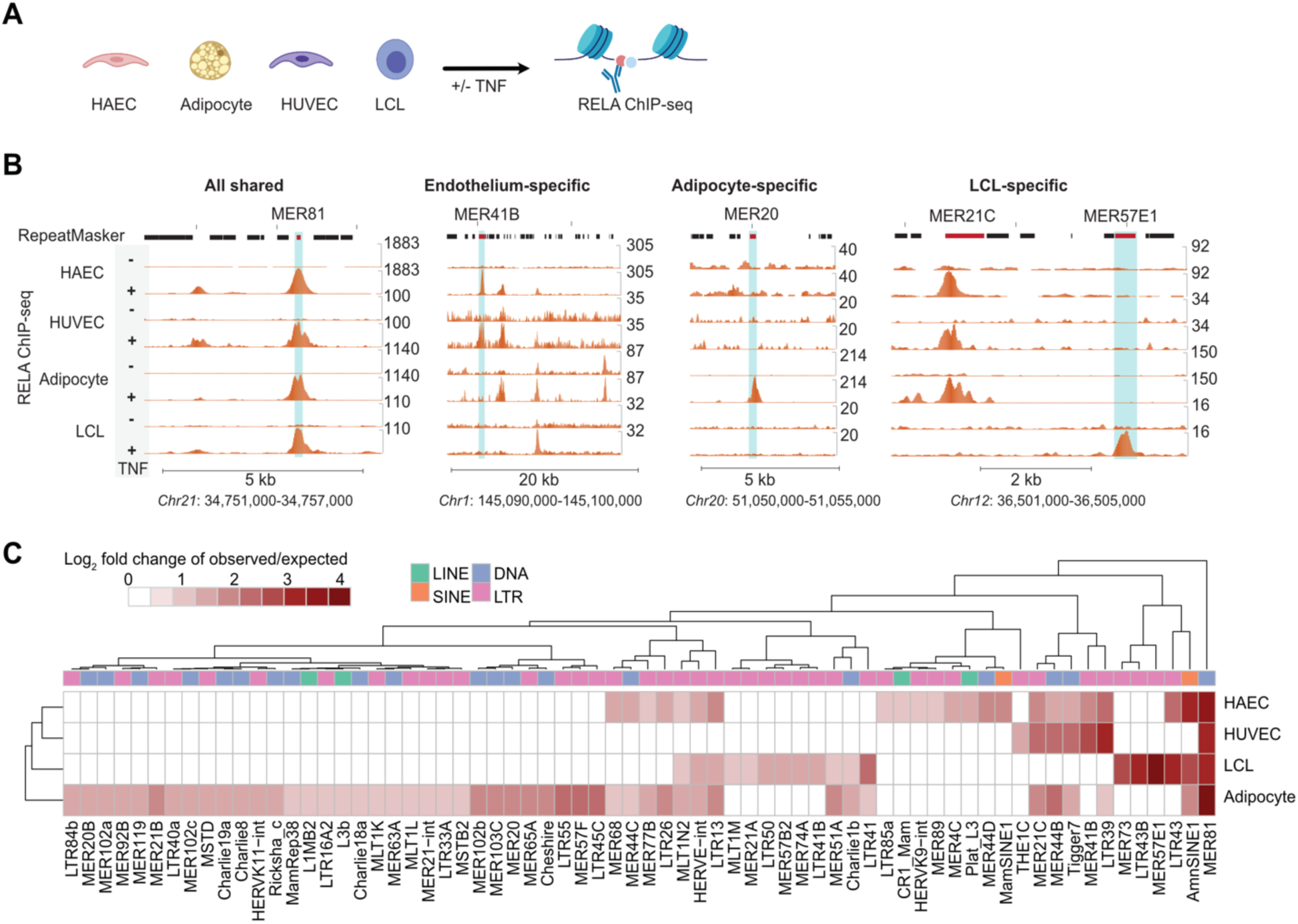
Multiple TE subfamilies were enriched within RELA-bound regions in different human cell types. A) A schematic showing the use of RELA ChIP-seq data from four human cell types. Together with RELA ChIP-seq data from this study, another three public RELA ChIP-seq datasets were incorporated to assess TE contribution to RELA-bound regions across cell types, which covers HAEC (Human Aortic Endothelial Cells), HUVEC (Human Umbilical Vein Endothelial Cells), LCL (Lymphoblastoid Cells) and Adipocytes. B) Examples of cell-type-shared and cell-type-specific TE-derived RELA-bound regions. C) An enrichment heatmap of identified TE subfamilies in the four cell types (cutoffs: FDR <0.05, log_2_ fold change > 1 and observed overlap events >= 15).

To gain further insight into the nature of the shared and cell-specific TE-derived RELA binding we investigated TF motif usage within TE-derived RELA-bound regions from each species and cell type individually. Supporting a direct role of DNA sequence features in NF-κB recruitment, motifs from RELA and other NF-κB subunits (RELB, REL, NFKB1, and NFKB2) were consistently detected in most TE subfamilies from all the species and cell types. For example, 16 out of 22 subfamilies in HAEC showed strong NF-κB motif enrichments (Figure 5A, 5B and Supplementary Figure 6). In endothelial cells, NF-κB collaborates with AP-1 for inducible transcriptional regulation (Oeckinghaus et al. 2011; Dorrington and Fraser 2019).

**Figure 5.**
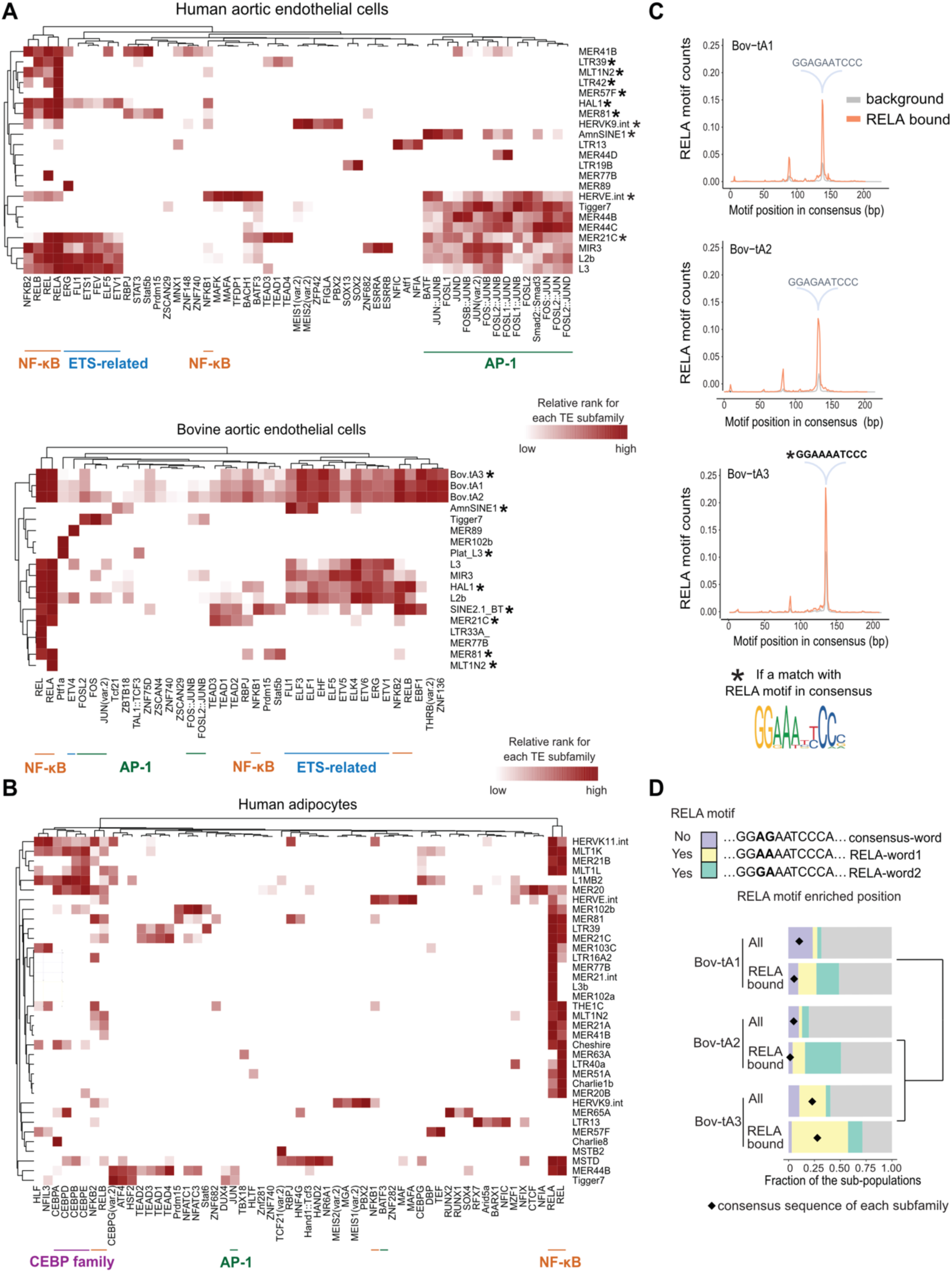
Multiple TF motifs contribute to TE-derived RELA bound regions across species and cell types. Heatmaps of significantly enriched TF motifs from A) human aortic endothelial cells as well as cow aortic endothelial cells, and B) human adipocytes. Significantly enriched TE subfamilies from species/ cell type analyses were investigated and only TEs detected with significant TF motifs are shown in the heatmaps. Rows and columns of the heatmaps indicate TE subfamilies and TF motifs, respectively. The intensity of heatmaps represent the rank of significantly enriched TF motifs for each TE (see more details in Methods). TF members of NF-κB, ETS, AP-1 and adipocyte-specific factors have been highlighted. C) RELA motif distribution relative to the consensuses of Bov-tA1, Bov-tA2 and Bov-tA3. Gray and orange represent all instances and RELA-bound instances, respectively. D) Fine stratification of Bov-tA1/2/3 instances based on abundant RELA motif-words. The left panel shows three investigated motif-words, including a consensus word of Bov-tA1/2 (consensus-word), and the top two abundant RELA motif-words (RELA-word1 and RELA-word2). Note RELA-word1 is the consensus motif of Bov-tA3. The right panel displays the fraction of instances that match with each word for all instances and RELA-bound instances. Purple, yellow, green and gray represent consensus-word, RELA-word1, RELA-word2 and all other sequences, respectively. The dendrogram depicts the phylogenetic relationship of Bov-tA1/2/3, estimated by the similarity of their consensus sequences.

Accordingly, we found enrichments for motifs from AP-1 subunits (e.g., JUN and FOS) in specific TE subfamilies enriched in RELA binding data from HAEC, HUVEC and cow aortic endothelial cells (Figure 5A and Supplementary Figure 6B). We also found multiple transposable elements with high AP-1 motif enrichment that lacked NF-κB motif enrichment (such as Tigger7 and MER44B/C/D in HAEC). This suggests that a subset of TE-derived RELA bound elements arise from co-binding of NF-κB and AP-1 to non-canonical/degenerate NF-κB sites (Kolovos et al. 2016). Additional evidence for the collaboration of ETS-related factors in TE-derived RELA bound regions was observed in aortic endothelial cells from human and cow (7/22 and 11/18 enriched TE subfamilies in human and cow, respectively). This finding could be driven by ERG binding, a known ETS factor that plays a critical role in endothelial function and inflammation (Shah et al. 2016; Hogan et al. 2017). Further supporting the importance of collaborative TF binding in TE-derived RELA bound regions, we found enrichments of 9 subfamilies (e.g., MER20 and L1MB2) for the adipocyte-specific master regulator CEBPA in human adipocyte RELA ChIP-seq data (Freytag et al. 1994; Darlington et al. 1998) (Figure 5B). These results indicate that further cell-type specific ChIP-seq experiments are needed to capture the full repertoire of TE subfamilies associated with NF-κB binding.

### Bovine SINEs mediate RELA binding via a restricted repertoire of RELA motifs

To further investigate the role of TF binding motifs in TE-derived RELA bound regions we investigated the sequence features within the ∼14,000 RELA-bound regions observed in the bovine SINE subfamilies Bov-tA1, Bov-tA2, and Bov-tA3. These three bovine SINE subfamilies make up ∼15% of all RELA binding observed in bovine aortic endothelial cells. Bov-tA elements originated through recombination between tRNA-Glu derived SINEs and a fragment of BovB element before the radiation of Pecoran ruminants (∼25-30 MYA) (Nijman et al. 2002; Nilsson et al. 2012). Collectively, Bov-tA1/2/3 give rise to more than 750,000 copies in the modern cow genome with a consensus sequence length of ∼210 bp. Given the sequence divergence of transposon instances relative to the subfamily consensus, Bov-tA1 are estimated to be the oldest subfamily, followed by Bov-tA2 with Bov-tA3 being the youngest one (Figure 2C). Unlike the Bov-tA1 and Bov-tA2 consensus sequences, the Bov-tA3 consensus sequence contains a RELA motif. We found that a one nucleotide difference at the 130-140 bp position of the aligned consensus sequences can explain the presence of the RELA motif in Bov-tA3 (Supplementary Figure 7A).

To investigate how the presence of a RELA motif relates to RELA binding observed by ChIP-seq, we compared RELA motif density of RELA-bound Bov-tA1, Bov-tA2, and Bov-tA3 instances to unbound instances of the same repeat subfamily. Indeed, we observed a significant increase in motif density at the 130-140 bp position in RELA bound regions (chi-squared test; 4.4-fold, 3.8-fold, 2.0-fold enrichment for Bov-tA1/2/3, respectively; P-value<2.2×10^-16^ for all three subfamilies; Figure 5C). Given that a single mutation at the 130-140 bp position of Bov-tA1 or Bov-tA2 can generate a RELA motif, which in turn could be spread through the genome, we next asked if the RELA bound instances were biased towards specific motif words. We tabulated all 10 bp “words” in the 130-140 bp region that matched the RELA motif. The most abundant RELA motif matching word (“RELA-word1”) corresponded to the consensus sequence of Bov-tA3 and accounted for 18%, 12%, and 54% of RELA-bound Bov-tA1, Bov-tA2, and Bov-tA3 elements, respectively (Supplementary Figure 7B). The second most frequent RELA motif containing word (“RELA-word2”) was also found in all three subfamilies and accounted for 22%, 35%, and 14% of RELA-bound Bov-tA1, Bov-tA2, and Bov-tA3 elements, respectively. The presence of “RELA-word1” and “RELA-word2” in RELA-bound instances was significantly above background expectations for all three subfamilies (Supplementary Figure 7B; chi-squared test; p-value <2.2×10^-16^ for all three subfamilies; Figure 5D). Overall this analysis shows that while the consensus sequences for Bov-tA1 and Bov-tA2 do not possess a canonical RELA motif, a single base pair mutation can create one for RELA recruitment in Bov-tA1/2 instances.

We also observed an enrichment of RELA motifs at a specific position (378-387 bp) of the consensus sequence and an increased fraction of RELA motif-word-matched subpopulations in RELA-bound instances in MER41B in human aortic endothelial cells (Supplementary Figure 7C and 7D). The other potential example of this phenomenon is LTR13, which lacks a RELA motif in the consensus but enriches for RELA binding. However, we did not find evidence for a site-specific RELA motif within this subfamily. While direct experimental evidence is needed, it is intriguing to speculate that the presence of RELA motif-containing instances of these subfamilies played a role in their expansion and evolution in mammals.

### DNA transposon MER81 carries two RELA motifs responsible for enhancer activity

MER81 was the only TE subfamily enriched in RELA bound regions from all tested species and cell types. Using RepeatMasker annotations from a representative set of 24 mammals (Chuong *et al*. 2016), we found MER81 to be present in all 23 placental mammals and absent in the non-placental mammal representative opossum (Supplementary Table 1). In placental mammals, a median of ∼2800 copies were identified in each species. There is a notable lineage-specific reduction of annotated MER81 elements in Eulipotyphla (hedgehog) and Rodentia (mouse, rat and prairie vole) species (∼500 copies in each species). In human, 3639 MER81 elements were annotated in the genome, among which 734 had RELA motifs. From our HAEC ChIP-seq data, we identified 87 extant MER81 copies bound by RELA in response to TNF stimulation.

Consistent with the general features of RELA-bound TEs, RELA-bound MER81s also exhibit active CRE characteristics measured by H3K4me2, H3K27ac and ATAC-seq signal before and after TNF stimulation (Figure 6A, Supplementary Figure 8A). Furthermore, RELA-bound MER81 elements were enriched in the vicinity of genes that are annotated with immune-related functions (Supplementary Figure 8B), such as “increased hematopoietic cell number” (binomial p value= 4.4×10^-6^) and “IFN-gamma pathway” (binomial p value= 5.5×10^-5^). In particular, we observed the association with multiple immune/inflammatory genes. For example, the proximal promoter region (0.3kb upstream of the TSS) of the gene *CXCR5*, which plays an important role in B cell maturation and migration (Pereira et al. 2010), has one instance of MER81 element with RELA binding that is shared across all four human cell types. We also found several interferon-signaling related genes in the vicinity of RELA-bound MER81 instances, including *IFNAR1* (surface receptor of IFN-α signaling; 58 kb downstream of the TSS), *TYK2* (downstream kinase required for IFN-α signaling; 5 kb upstream of the TSS), *IFNGR2* (surface receptor of IFN-γ signaling; 19 kb upstream of the TSS), *JAK2* (downstream kinase required for IFN-γ signaling; 47 kb upstream of the TSS), *IRF1* (known transcription factor activated by IFN-γ signaling; 15 kb downstream of the TSS) (Supplementary Table 2).

**Figure 6.**
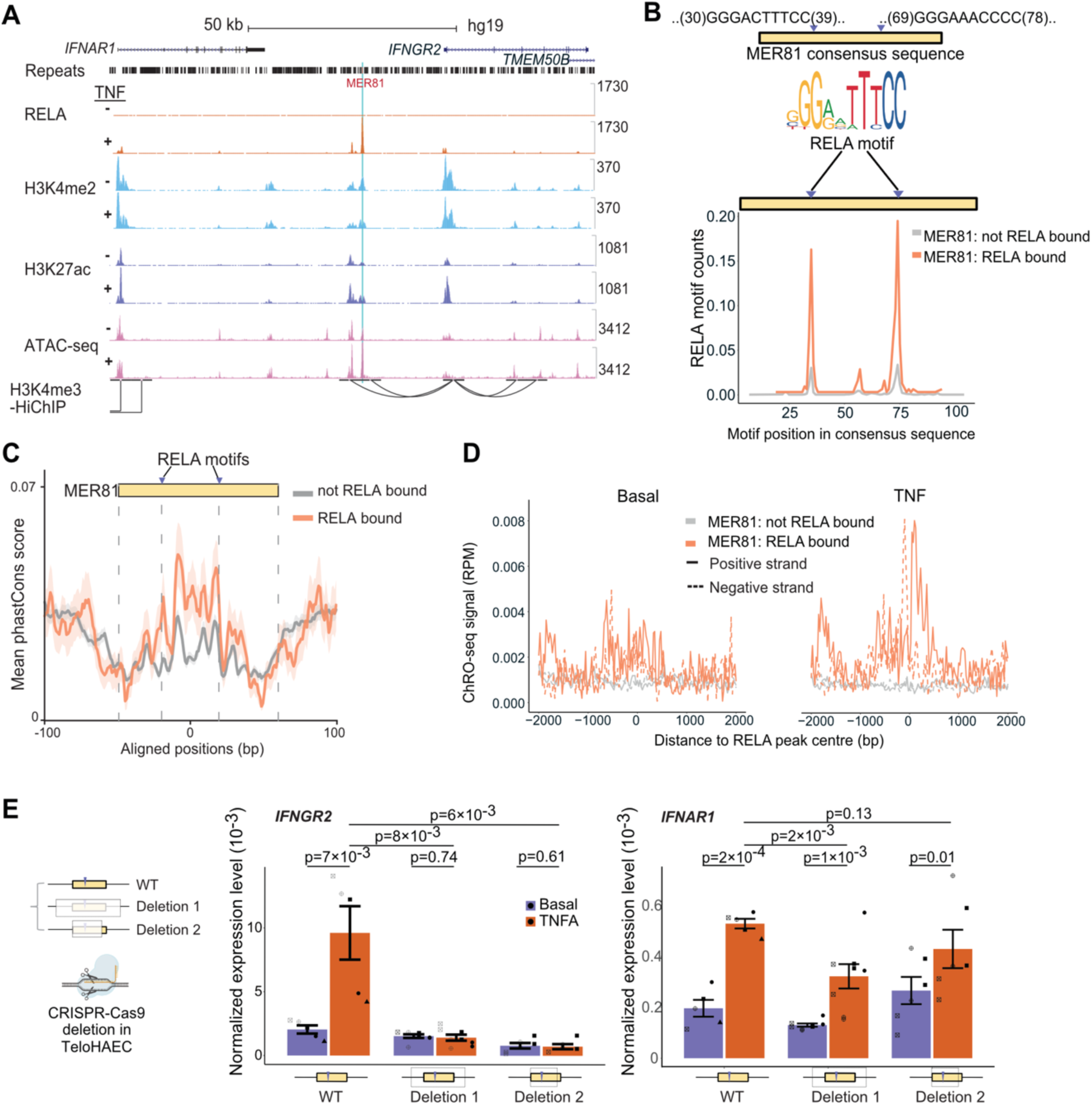
DNA transposon MER81 carries two RELA motifs responsible for enhancer activity. A) An instance of a MER81-derived RELA-bound region. The genome browser view covers tracks of RELA, H3K27ac ChIP-seq, ATAC-seq and H3K4me3 HiChIP. “-” and “+” denote basal and TNF stimulation, respectively. B) Two canonical RELA motifs in the MER81 consensus sequence appear to function for RELA binding. The left panel shows the two RELA motifs in the consensus sequence, while the right panel depicts normalized motif counts of RELA motifs in extant human MER81 instances. Orange and gray represent RELA-bound and no RELA-bound instances, respectively. C) Aligned conservation scores around RELA-bound vs no RELA-bound MER81 instances. All the annotated MER81 instances were first aligned to their consensus sequence and then calculated for the mean phastCons score for each aligned position. D) eRNA signal around MER81-derived RELA sites before and after TNF stimulation. eRNA signal was obtained from TeloHAEC ChRO-seq data. E) Functional validation of the regulatory activity of a native MER81 sequence. This selected MER81 instance was deleted with different precisions (Deletion 1/2) using *CRISPR-Cas9* in TeloHAECs. The plot shows the qPCR-measured expression levels of *IFNGR2* and *IFNAR1* in wild-type and two MER81 knockout TeloHAEC lines (with/without TNF simulation for 45 min). For E), we have performed paired one-sided *t*-test for comparisons that involve the same genotype (e.g., stimulated vs unstimulated mutants) and unpaired one-sided *t*-test for ones that involve different genotypes (e.g., TNF stimulated wild-type vs Deletion 1). Error bars indicate standard error of the mean. Each biological replicate is shown with a unique shape in the figure.

To understand how MER81 could recruit RELA for binding, we examined its sequence features. We found two canonical RELA motifs in its 114-bp consensus sequence, which could explain RELA binding (Figure 6B). To expand the repertoire of RELA-bound MER81 elements, we additionally searched the pool of annotated RELA-bound regions from the GTRD database, which contains data from several human cell lines (Supplementary Material 2). Together with our HAEC data, we obtained 327 MER81 instances that overlapped a RELA peak summit in at least one human cell type. We observed a clear enrichment of normalized RELA motif counts in RELA-bound MER81 elements compared to unbound ones (0.16 vs 0.03 for MER81 RELA-motif 1 and 0.19 vs 0.03 for MER81 RELA-motif 2) (Figure 6B). In other words, the loss of RELA binding in extant MER81 copies could be largely explained by the loss of RELA motifs. In addition, RELA-bound MER81 sequences show increased sequence constraints relative to unbound ones (34% increase in averaged phastCons score, one-sided *t*-test, p-value=0.008), further indicating their potential functional importance (Figure 6C).

Further support that these RELA-bound MER81 sequences can serve as enhancers, was obtained by ChRO-seq analysis revealing the bi-directional signal associated with active enhancers which are modulated by TNF stimulation. This feature was not seen at MER81 elements that were not bound by RELA (Figure 6D). To further explore the functionality of these MER81 elements, we performed luciferase reporter assays to assess the enhancer activity of the sequences. We first tested the activity of the MER81 consensus sequence, which serves as an “ancestral” representative of all MER81 instances. We found a modest increase enhancer activity of the MER81 consensus sequence compared to an empty vector control under basal condition (one-sided t-test; 1.6-fold, p-value=0.03), which was enhanced upon TNF stimulation (one-sided t-test; 3.0-fold, p-value=0.02). This modest enhancer activity of the putative ancestral MER81 sequence was lost when the RELA motifs within the sequence were disrupted (one-sided t-test; p-values of 0.81, 0.10 and 0.40 for TNF-stimulated mutant1, mutant2, and mutant3, respectively; Figure 6E and Supplementary Figure 9A). Altogether, our results are consistent with our ChIP-seq results showing the MER81 instances containing intact RELA motifs can enhance gene expression after TNF stimulation.

Next, we selected one native RELA-bound MER81 element in human that resides in the vicinity of interferon signaling genes to explore its potential enhancer function using the same luciferase assay (Figure 6A). This instance is located between two interferon receptor genes, *IFNAR1* and *IFNGR2* (+58 kb and -19 kb relative to *IFNAR1* and *IFNGR2* TSS, respectively), both of which we identified as RELA target genes in our HAEC RNA-seq data. Our epigenomic data indicate that this MER81 sequence coincides with the largest RELA peak at the locus and is within an accessible chromatin region with a strong H3K4me2/H3K27ac signal. Using our H3K4me3 HiChIP data from TeloHAECs we identified a chromatin loop connecting this instance and the downstream target gene *IFNGR2*. We first attempted to validate the enhancer activity of this sequence via luciferase reporter assay (Supplementary Figure 9B). This instance only retained the first RELA motif. Consistent with our luciferase results using the ancestral MER81 sequence with a single RELA motif showed little to no enhancer activity which increased slightly in response to TNF stimulation (one-sided t-test for TNF-stimulated native sequence, 1.5-fold, p-value=0.07) whereas the motif-shuffled counterpart exhibited loss of enhancer activity (one-sided t-test for TNF-stimulated mutant sequence, 0.91-fold, p-value = 0.73; Supplementary Figure 9B).

To test the function of this enhancer in its chromatin context, we deleted the genomic region in TeloHAECs using *CRISPR-Cas9* (Supplementary Figure 9C). We performed two deletion experiments: “Deletion 1” which spans from 160 bp upstream to 163 bp downstream of the 101-bp MER81 instance; and “Deletion 2” which is restricted to the first ∼80 bp of this MER81 instance and includes the RELA motif. We then checked the effects of these deletions on the neighboring *IFNAR1* and *IFNGR2* genes (note *IFNGR2* had an order of magnitude higher expression than *IFNAR1* before and after TNF). Both *IFNAR1* and *IFNGR2* were significantly induced by TNF (2.7-fold change of pre-mRNA transcript expression; paired one-sided *t*-test, p-value of 2×10^-4^ and 4.7-fold change of pre-mRNA transcript expression; paired one-sided *t*-test, p-value of 7×10^-3^ respectively). We found a small but significant effect on induced expression in *IFNAR1* upon the loss of this MER81 instance in Deletion 1 (0.61-fold change; unpaired one-sided *t*-test, p-value of 2×10^-3^) and no significant change in the more specific Deletion 2 (0.81-fold change; unpaired one-sided *t-*test, 0.13; Figure 6F). In contrast, both Deletion 1 and Deletion 2 abolished the ability of *IFNGR2* to be induced by TNF (paired one-sided t-test, p-value of 0.74 and 0.61 for Deletion 1 and 2, respectively; Figure 6F). These results indicate that this MER81 instance functions as a *bona fide* enhancer, which is required for the regulation of the downstream target gene *IFNGR2*. Furthermore, the fine-mapping of the functional sequence (Deletion 2) reveals an essential 80 bp fragment in this element which encompasses one intact ancestral RELA motif. Altogether, our results support that MER81 has functional RELA motifs capable of regulating target gene expression *in vivo*.

### Ancestral reconstruction reveals a stabilization of RELA motifs during MER81 evolution

The enrichment of RELA binding in extant MER81 copies appears to be a common feature across cells and mammalian lineages. However, the extant copies of MER81 with RELA binding ability tend to be species-specific. This is underscored by the mouse-specific loss of MER81 copies, with only 500 elements remaining in the mouse genome, compared with ∼3600 in human and ∼2600 in cow. Furthermore, 79% of MER81 derived RELA binding events in human are human-specific (Supplementary Material 3). To delineate the evolutionary trajectories of the MER81 elements over the course of human evolution, we utilized ancestral whole-genome reconstructions of human and ten other mammalian species which cover an evolutionary time window of ∼100 million years (Campitelli et al. 2022). With this strategy, we were able to recover the genome sequences of each node/common ancestor (Figure 7A). By pooling all ten reconstructed genomes together with the modern human genome, we achieved 11 nodes of genome sequences and were able to study mammalian and primate evolution from a human-centric perspective. We found the total copy number of MER81 was highly stable during evolution; in particular, ∼2800 MER81 elements were detected in each node. The stability of MER81 copies suggests no or a low level of transposition activity in the recent ∼100 million years. Secondly, a large proportion (20-45%) of MER81 elements were identified with RELA motifs and the proportion containing a RELA motif decreased steadily over time, with >45% of MER81 elements containing a RELA motif in node 10 (or the human-star-nosed-mole common ancestor) to only ∼20% containing a RELA motif in modern human (Figure 7B).

**Figure 7.**
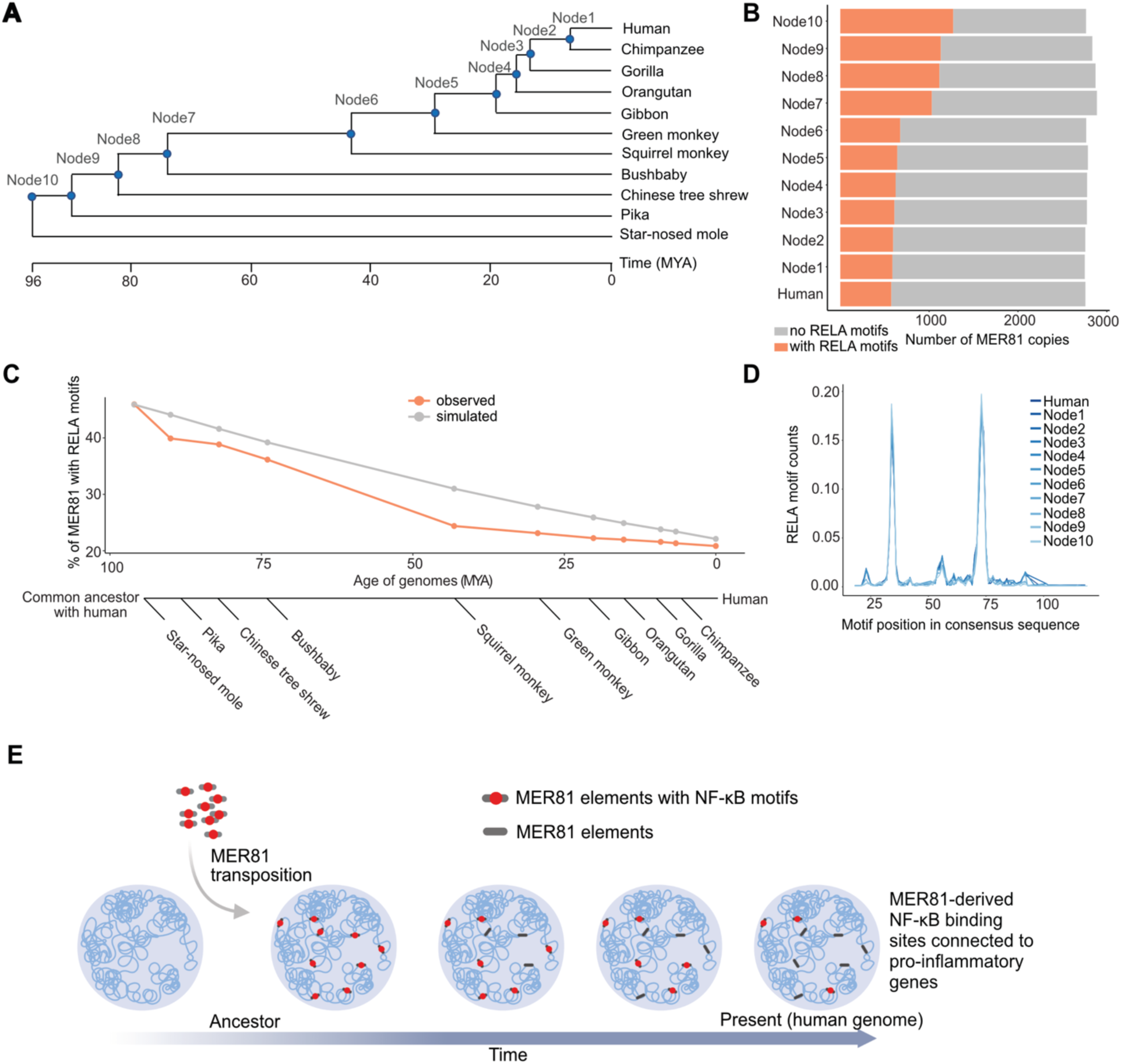
Ancestral reconstruction reveals a stabilization of RELA motifs during MER81 evolution. Modern human together with ten related mammalian species were selected to reconstruct a series of ancestral genomes based on multiple sequence alignments. A) Phylogenetic tree of the 11 selected species to reconstruct ancestral genomes. Each node in the tree represents a reconstructed genome of the corresponding common ancestor. All the nodes were collected to form a series of representative time points in the process of human evolution. The phylogenetic tree was obtained by querying TIMETREE database (Kumar et al. 2017). B) The number of MER81 copies in each human ancestor genome. Gray and orange represent total numbers and copies containing RELA motifs, respectively. C) RELA motif change in MER81 along human evolution. The percentage of MER81 proportion containing RELA motifs is shown as a function of the estimated evolutionary age of each node/common ancestor (divergence time estimation obtained from TIMETREE database (Kumar et al. 2017)). A simulation of MER81 sequence change based on nucleotide substitution was shown in gray (see Methods). D) The distribution of RELA motifs in MER81 instances in all human ancestors. Only instances with detected RELA motifs were investigated. E) Schematic of a MER81 co-option model during human evolution. MER81 colonized into the genome at a very early stage. These elements were then subject to a long-term selection with no or low level of expansion by active transposition. Functional units were potentially co-opted as enhancers to regulate target gene expression. Panel E was created with BioRender.com.

Relative to a simulated RELA motif decay rate derived using the estimated nucleotide substitution rate (see Methods), we observed a rapid decrease of RELA motifs in the early stages of MER81 colonization followed by a stabilizing trend starting around the last common ancestor of human and squirrel monkey (Figure 7C). To put this evolutionary trajectory into context, we expanded our analysis to include the additional human TE subfamilies associated with RELA binding. We also queried the motif for the AP-1 family transcription factor JUN, which is well known to bind together with NF-κB, and is prominently expressed in endothelial cells (Oeckinghaus et al. 2011; Dorrington and Fraser 2019). We also found JUN motifs to be enriched within distinct subsets of TE-associated RELA binding sites (Figure 5). While we observed no signal from scrambled RELA or JUN motifs, distinct deceleration patterns of RELA and JUN motif decay from multiple transposons were observed (e.g. MER57F, MER77B and MER44D; Supplementary Figure 10). The high and stable signals of JUN motifs, and the absence of RELA motif signal, from MER44B/C/D is also consistent with our motif enrichment analysis (Figure 5A), as well the motif enrichment analyses obtained from accessible chromatin regions induced by the bacterial infection of macrophages (Bogdan et al. 2020). By contrast, the NF-κB binding in MER81 (which was the only significant TF motif we found enriched in this subfamily; Figure 5A) showed no signal for JUN motifs.

We further investigated the distribution of RELA motifs within MER81 elements that contain the motif, and confirmed that these motifs were still centered on the two positions from the consensus sequence and that no apparent motif turnover or shift occurred within these elements (Figure 7D). Together, our analysis supports a model where the DNA transposon MER81 colonized the genome prior to the radiation of placental mammals, followed by a long-term selection for the NF-κB motifs it contained (Figure 7E). At least a subset of these elements may have been co-opted as active enhancers to regulate target gene expression in an NF-κB-dependent manner.

## DISCUSSION

In this study, we explored the contribution of transposable elements to genome-wide binding profiles of NF-κB in three mammalian lineages. Our comparative analyses of RELA binding, together with epigenetic and transcriptional profiles, revealed conserved and lineage-specific TE subfamilies that are likely to participate in mammalian NF-κB regulatory networks. Given the active enhancer features and proximity to multiple NF-κB target genes, it is likely that a subset of these TE-derived RELA-bound elements have been incorporated into the host NF-κB regulatory networks.

The expansion of RELA-bound regions in the bovine genome largely results from Bovine SINE subfamilies Bov-tA1/2/3, which contribute over 14,000 RELA-bound regions to the cow genome (Figure 2A, 2B). This bovine-specific expansion even exceeds the ∼10,000 SINE-B2 derived CTCF sites observed in the mouse genome (Bourque et al. 2008; Schmidt et al. 2012), making it one of the most striking examples of lineage-specific TF binding expansions described in mammals to date.

While the null hypothesis is that the vast majority of the Bov-tA1/2/3 derived NF-κB sites would be non-functional, the large number of these lineage-specific expansion of Bov-tA1/2/3 raises the possibility that a small subset of these elements influences ruminant gene regulatory networks. Kelly *et al*. 2022 recently discovered a substantial expansion of IFNγ-inducible elements by a ruminant-specific SINE, Bov-A2 (∼12,000 instances), for the regulatory evolution of ruminant innate immunity. Bov-A2 originated from Bov-tA elements via dimerization of the Bov-tA tail regions after the radiation of Pecoran ruminants (>10 million years ago) (Nijman et al. 2002; Nilsson et al. 2012; Bao et al. 2015). Bov-A2 and Bov-tA1/2/3 constitute the family BovA, and the related evolutionary trajectory highlights a potentially pervasive function of the BovA family in ruminant innate immunity. For both Bov-tA1 and Bov-tA2, we observed evidence of “maturation” of transposon instances from imperfect “pre-sites” or “proto-motifs” in ancestral sequences to perfectly matched RELA binding motifs. This phenomenon, currently described as the “proto-motif” model (MacArthur and Brookfield 2004; Bourque et al. 2008; Zemojtel et al. 2011; Sundaram and Wysocka 2020; Judd et al. 2021), involves individual TEs acquiring their TF-binding motifs after insertion. Such a process could give rise to a Bov-tA1 subfamily heterogenous for their ability to interact with NF-κB. Subsequently, the ancestral Bov-tA2 element, which also lacks an ancestral RELA binding motif, could have independently evolved and propagated the same two common RELA motif-words. Finally, the ancestral sequence of the Bov-tA3 element, which contains the most prevalent RELA-motif word, contributed disproportionately to the observed Bov-tA derived RELA binding events (Figure 2B). Furthermore, Bov-tA3 also shows evidence of expansions of RELA-word2 supporting a model where the gain and loss of RELA motifs was an ongoing process for Bov-tA elements during the evolution of ruminants. In the discussion of their study of *Alu* associated NF-κB sites, Antonaki et al. proposed that such sites could serve as transcriptionally inert docking sites of NF-κB which could prevent excessive targeting of promoters (Antonaki et al. 2011). Likewise, one could speculate that the large number of Bov-tA associated NF-κB binding sites could be tolerated if they somehow collectively provide a fitness advantage by attenuating the bovine inflammatory response.

It is intriguing to speculate that the Bov-tA (and other TE subfamilies) expanded their foothold in the genome by promoting their own expression through NF-κB binding. While studies have demonstrated NF-κB activation in relevant cell types and developmental stages in a variety of mammals (e.g. bovine oocytes and preimplantation embryos (Paciolla et al. 2011) as well as rat and mouse testes (Delfino and Walker 1998) (Rasoulpour and Boekelheide 2005) (Lilienbaum et al. 2000), no experimental evidence exists to support such a mechanism.

Prior work demonstrated the ability of *Alu* sequences to bind RELA and suggested that *Alu* elements contributed substantially to RELA binding in the human genome (Apostolou and Thanos 2008; Antonaki et al. 2011). While our study in primary endothelial cells did not identify *Alu* as a major contributor of TE derived RELA bound regions, it is important to note that the reported NF-κB motif containing *Alu* transposons (including AluSx, AluSg and AluY) are young elements (Bennett et al. 2008), many of which would not be uniquely mappable with the 100 bp single end reads used in our RELA ChIP-seq experiments (Sexton and Han 2019). Another explanation for this discrepancy with the prior work comes from our stringent definition for TE derived RELA bound regions, which required an overlap between a known TE and the summit of RELA peak. In comparison, Antonaki et al. 2011 allowed for an overlap with any portion of a RELA peak.

Growing evidence has indicated that TEs, especially evolutionarily young TEs, act as a source of *cis*-regulatory elements, which primarily innovate regulatory programs in a species-and tissue-specific manner (Buecker and Wysocka 2012; Chuong et al. 2016; Fuentes et al. 2018; Todd et al. 2019; Du et al. 2022; Kelly et al. 2022; Lee et al. 2022; Choudhary et al. 2023).

Although fewer in number, older TEs have been demonstrated to contribute to species-specific (Notwell et al. 2015) or conserved (Lynch et al. 2011) gene regulatory networks. Our comparative epigenomic study of TE-derived RELA binding allowed us to identify 24 TE subfamilies enriched within RELA-bound regions which appear to predate the human-mouse and mouse-cow divergence times (Figure 2C). Of these TE subfamilies, AmnSINE1, UCON26, Plat_L3, MamSINE1, MER21C and MER81 were found to be enriched for RELA binding in at least two species. Given the age of these elements it is quite possible that additional instances may exist that could not be readily identified by RepeatMasker. Nonetheless, the ∼300 million-year-old AmnSINE1 subfamily has been shown to contribute TF binding sites to mammalian gene regulatory networks (Sasaki et al. 2008; Hirakawa et al. 2009; Chuong et al. 2016) and our results suggest AmnSINE1 derived NF-κB binding may contribute its enhancer function across species and cell types. Much less is known about the LTR retrotransposon MER21C aside from work demonstrating this element formed part of the promoter of the apoptosis inhibitory *NAIP* gene in humans (Romanish et al. 2007). While it is unlikely these examples of TE associated RELA binding shared across species are due to systematic errors in TE annotation, it is important to acknowledge that a recent study estimated that 10-14% of *Alu* and L1 subfamilies are discordantly annotated in human and chimpanzee genomes (Carey et al. 2021).

One of the central discoveries in this study is the contribution of the ancient DNA transposon, MER81, to NF-κB mediated regulatory networks. Most MER81-derived RELA-bound regions are species-specific (79%) indicating the ancient MER81 elements may play core NF-κB functions in a human-specific manner. In human, the majority (>80%) of MER81-derived RELA-bound regions are shared across cell types, which increases the potential for MER81 to influence core components of NF-κB mediated transcriptional regulation in different cellular contexts. This observation is supported by a recent study that found that: 1) the chromatin accessibility of MER81 elements increased after bacterial or viral infection of human macrophages; and 2) that these elements showed unique enrichment for NF-κB motifs (Chen et al. 2023). We demonstrated that one MER81 instance can control the activation of the IFN-γ signaling gene *IFNGR2* (Figure 6A and 6F). It is possible that RELA bound MER81 elements play a larger functional role in IFN-γ signaling, given their significant enrichment near additional IFN-γ signaling pathway genes (seven genes in addition to *IFNGR2*).

Our study presents evidence highlighting the significance of the MER81 derived RELA motif(s) in promoting NF-κB binding. However, it is important to acknowledge that other chromatin features and transcription factor (TF) motifs, not necessarily originating from transposable elements (TEs), likely play a role in enabling this TE to function as a CRE in vivo. For example, it has been well established that regions robustly bound by NF-κB typically contain multiple high affinity motifs within an accessible chromatin context (e.g., (Michida et al. 2020; Alizada et al. 2021). This point was also made in this study considering the modest ability of the MER81 element at the human *IFNGR2* locus to increase gene expression in a luciferase reporter assay in response to TNF. While we only found enrichments for NF-κB motifs in MER81 elements, we did find other RABS with clear enrichments for the motifs of TFs prominently expressed in endothelial cells (namely AP-1 and ETS family members). For example, several AP-1 enriched TE subfamilies were found in the MER44B/C/D and Tigger subfamilies, all of which lacked the enrichment of canonical NF-κB motifs. This observation is consistent with previous work showing the co-binding of AP-1 and NF-κB at regions with degenerate NF-κB motifs (Kolovos et al. 2016). Notably, both AP-1 and NF-κB motifs were also found to be significantly enriched within several TE subfamilies (such as MER41B and THE1C) in experiments performed on pathogen stimulated macrophages (Bogdan et al. 2020). While additional higher throughput experiments focusing on the collaborating TFs and chromatin features are needed, our results demonstrate the ability of TE-derived NF-κB motifs to contribute to RELA binding at active CREs.

TF binding sites provided by ancient TEs can be thought of as “natural experiments” that can be followed through speciation events and studied in situ in relevant cell types. For example MER81, an ancient DNA transposon with low replicative efficiency spread a short DNA sequence containing two RELA motifs, via a “cut-and-paste” mechanism prior to the divergence of placental mammals (Muñoz-López and García-Pérez 2010; Wells and Feschotte 2020). By using ancestral genomes to trace the RELA motif instances in MER81 (and other TEs subfamilies) during the last ∼100 million years of mammalian evolution leading to the human lineage, we observed a pattern that could indicate a short phase of negative selection against a subset of functional MER81 elements, followed by positive selection of another subset of these elements that now play a functional role in NF-κB regulatory networks (Figure 7C). While almost all these TE-associated RELA binding sites await functional validation, our work provides additional evidence supporting the recurrent co-option of TEs into NF-κB regulatory networks during mammalian evolution.

## METHODS

### ChIP-seq

To annotate mouse TE elements we performed H3K4me2 and H3K27ac ChIP-seq with and without TNF treatment from mouse aortic endothelial cells. One biological replicate of primary mouse aortic endothelial cells (Cell Biologics, cat# C57-6052, lot# B092913T2MP) was selected and cultured following the methods of Alizada et al. 2021. Cells were then treated with 10 ng/mL recombinant mouse TNF (Cell Applications, cat# RP2031-20) for 45 min in basal Endothelial Cell Growth Media MV2 (PromoCell) without supplements. The unstimulated control samples were treated with vehicles (i.e., an equivalent amount of water in MV2 media without supplements). For the ChIP-seq experiments, the primary aortic ECs were starved (basal media without supplements) for 16 h prior to TNF stimulation. Next, cells were crosslinked in 1% formaldehyde for 10 min at room temperature and were then lysed for nuclei isolation as described in Schmidt *et al*., 2009 (Schmidt et al. 2009). Chromatin was sheared into 100–500-bp DNA fragments by sonication (Misonix Sonicator). ChIPs were performed at 4 °C overnight with 10 μg of antibody in a final volume of 250-μl block solution (0.5% BSA (w/v) in PBS): mouse anti-H3K27ac monoclonal (Millipore # 05-1334), rabbit anti-H3K4me2 polyclonal (Millipore # 07-030). The amplified and barcoded library was then selected for 200–350-bp fragments using Pippin Prep (Sage Science) and sequenced on Illumina HiSeq 2500 with a 100-bp single-end run to obtain ∼20–25 million single-end reads per sample.

### H3K4me3 based HiChIP

We prepared TeloHAECs with/without TNF stimulation for HiChIP experiments (two replicates per condition). Briefly, TeloHAECs were cultured following the protocol from Alizada et al. 2021. For TNF and vehicle treatment, the protocol was as described at the beginning of the ChIP-seq method, except here we used 10-ng/mL recombinant human TNF (Cell Applications, cat# RP1111-50). HiChIP was performed from a modified PLAC-seq protocol (Fang et al. 2016) in three steps as follows: **(1)** In situ proximity ligation. 5 million TeloHAECs per replicate were crosslinked in 37% formaldehyde as per instructions of the Arima Hi-C kit (cat# A510008) and frozen down as 10^6^ cells per aliquot. **(2)** Chromatin immunoprecipitation (ChIP). We adapted the protocol from Fang et al. 2016 with slight modifications. After proximity ligation, the nuclei were spun down at 2,500 g for 5 min at 4°C and the supernatant was discarded. The nuclei were then resuspended in 130 μl RIPA buffer (10 mM Tris, pH 8.0, 140 mM NaCl, 1 mM EDTA, 1% Triton X-100, 0.1% SDS, 0.1% sodium deoxycholate) with proteinase inhibitors. The nuclei were lysed on ice for 10 min and then sonicated using Diagenode Bioruptor® Pico with the following setting: 10 Cycles of 30 Secs On/Off. After sonication, 370 ul of RIPA buffer was added to the samples before they were cleared by centrifugation at 20,000 G for 20 min and the supernatant was collected. The clear cell lysate was mixed with 500ul of antibody (2.5 μg of H3K4me3, Millipore cat #04-745) coated Dynabeads M-280 (Sheep anti-Rabbit Thermo Scientific cat #11201D). The biotin pull-down procedure was performed according to Swift Biosciences Accel-NGS 2S Plus DNA Library Kit (cat# 21024). An optimal number of PCR cycles were determined by qPCR using the KAPA library quantification kit (cat# KK4824). Libraries were checked on a Bioanalyzer for quality and then sequenced on a NovaSeq6000 S4 for PE150.

### Luciferase reporter assay

For the investigated sequences, we designed 4 MER81 consensus-related sequences and 2 sequences from a native MER81 instance (located at *Chr21*:34755496-34755596 of hg19). The consensus sequence of MER81 was obtained from Repbase (Bao et al. 2015) and the native MER81 sequence was downloaded from the UCSC Genome Browser for coordinates of *Chr21*:34755496-34755596 (hg19). The mutant sequences were designed by randomly shuffling the RELA motifs in consensus/native sequences. We included a synthetic sequence with tandem 5 × RELA motifs as a positive control and generally observed > 20-fold change with TNF stimulation compared to empty pGL3 control for all experiments (values of data points are mentioned in the legend). Overall, we prepared 6 biological replicates with 3 technical replicates each for each investigated sequence to ensure reproducibility (See Supplemental Figure S9 for more details).

### *CRISPR-Cas9* deletions

Cas9 targeting guide RNAs (gRNAs) were selected flanking the selected MER81 instance (*Chr21*:34755496-34755596 of hg19) for deletion. On and off-target specificity of the gRNAs was calculated as described in Doench et.al 2016 (Doench et al. 2016) and Hsu et al. 2013 (Hsu et al. 2013) respectively, to choose optimal guides. Guide RNA sequences can be found in Supplemental Table 3. Guide RNA plasmids were assembled with gRNA sequences using the protocol described by (Mali et al. 2013). Briefly, two partially complementary 61-bp oligos were annealed and extended using Phusion polymerase (New England Biolabs). The resulting 100-bp fragment was assembled into an AflII-linearized gRNA empty vector (Addgene, ID#41824) using the Infusion assembly protocol (TaKaRa Bio). The sequence of the resulting guide gRNA plasmid was confirmed by sequencing with either T7 or SP6 primers. TeloHAECs were transfected with 5 µg each of the 5′ gRNA, 3′ gRNA, and pCas9_GFP plasmids (Addgene, ID#44719; (Ding et al. 2013)) using the Neon Transfection System (Life Technologies). Forty-eight hours post-transfection, GFP-positive cells were isolated on a BD FACSAria. After recovery, single cells were seeded in 96-well plates and inspected for colonies after 1–2 weeks. Upon 60% confluency, the colonies were split into replicate 96-well plates. Genotyping of the deletions was performed by amplifying products internal to and surrounding the target deletion. All deleted clones identified from the initial screen were sequenced across the deletion to confirm deletion boundaries. Four homozygously deleted clones were selected for the following experiments (as shown in Supplementary Figure 10).

### RNA isolation and qPCR

TeloHAECs were treated with/without TNF for 45 min. Total RNA was purified from single wells of >85% confluent six-well plates using the RNeasy Plus Mini Kit (Qiagen), and an additional DNase I step was used to remove genomic DNA. RNA was reverse-transcribed with random primers using the high-capacity cDNA synthesis kit (Thermo Fisher Scientific). Primary *IFNAR1* and *IFNGR2* gene expression was detected by primers targeting the first and fourth intron, respectively. The standard curve method was used to calculate expression levels using TeloHAEC genomic DNA to generate the standard curves.

Levels of *TUBA1A* RNA were used as an internal control to normalize expression values. The pimer sequences for *IFNAR1*, *IFNGR2* and *TUBA1A* can be found in Supplemental Table 3.

### Massively parallel sequencing sequencing data processing

RELA ChIP-seq data (including all cross-species and cross-cell-type RELA ChIP-seq data) were mapped as described in Alizada et al. 2021, followed by an additional quality filter (MAPQ >=3) to retain high-quality uniquely mapping reads. We compared this filtering to what we would obtain using the “XT:A:U” flag in the SAM file. For each of the three species, all of our MAPQ filtered reads (MAPQ>=3) were contained within the uniquely mapping reads obtained by the “XT:A:U” flag in the SAM file, suggesting that MAPQ >=3 is a sufficiently stringent means to obtain uniquely mapping reads for our analysis. The histone ChIP-seq and ATAC-seq data were analyzed as described in Alizada et al. (without MAPQ filtering). H3K4me3 HiChIP was processed using the MAPS pipeline (Juric et al. 2019) with default parameters. Significant loops were called using MAPS with default parameters for each sample. Since chromatin loops are expected to stay largely unchanged within a rapid induction system (in our case, TNF treatment for 45 minutes), we merged the loops identified from basal and TNF treatment conditions to generate a union set of active chromatin loops (in total 19372) for TeloHAECs. These loops vastly map active enhancer-promoter or active promoter-promoter connections (see Figure 6A for an example).

### Enrichment analysis for TEs overlapping RELA-bound regions

We used the GAT tool (https://gat.readthedocs.io/en/latest/contents.html) to determine the potential overrepresentation of TEs within RELA-bound regions. To run the enrichment analysis, we used the peak summits of RELA peaks, RepeatMasker annotated TE subfamilies. We set the workspace (available genomic regions for simulation) as all non-random chromosomes. 10000 iterations were requested for empirical distribution estimation and other parameters were set as default. The TE annotation files (matched annotation for hg19, mm10 and btau6) by RepeatMasker were downloaded using the table browser under the UCSC Genome Browser (https://genome.ucsc.edu). Simple repeats were excluded from all the TE annotations. In brief, the GAT tool shuffles RELA peak summits across the given workspace for 10000 iterations to obtain the distribution of expected overlap numbers for each TE subfamily. Empirical p-value and enrichment/fold change values are calculated by combining both observed and expected overlap numbers. The cutoffs for significantly enriched subfamilies across species were: FDR <0.05, log_2_ fold change >1 and observed overlap >=10 events. For cross-cell-type analysis, slightly more stringent cutoffs were used to control potential false positives: FDR <0.05, log_2_ fold change >1 and observed overlap >=15 events. The information of overlapping TEs for each RELA peak summit from human/mouse/cow can be found in Supplementary Material 4.

### Estimation of evolutionary ages of TEs

Age was calculated for each TE instance using percent divergence of sequence (obtained from the aforementioned TE annotation files from UCSC Table browser) divided by estimated nucleotide substitution rates of the corresponding species from (Bulmer et al. 1991; Pace et al. 2008). The median value of all instances from the same subfamily was calculated as the estimation of the subfamily age. Two point-estimations of human-mouse and mouse-cow divergence time (Figure 2C) were acquired from TIMETREE database (Kumar et al. 2017). Human-mouse divergence time was later used to divide TEs into “young” and “old” categories based on whether the predicted age of TEs was younger or older than when humans and mice diverged (in particular, the two categories were discussed in Figure 3).

### Analysis of TE overlap with TF bound regions

We took advantage of all the processed public TF ChIP-seq data from GTRD (Kolmykov et al. 2021) (v2018.3) to investigate TE and TF connections globally. The bound regions of TFs documented in GTRD were generated by merging ChIP-seq peaks from diverse conditions and cell types. This feature represents a comprehensive repertoire of bound regions for each TF, which allows us to study the general relationship between TEs and TFs in an unbiased manner. Note the way GTRD controls for multiple mapping reads in TF ChIP-seq data is slightly different from what we did for the multi-species RELA ChIP-seq data: GTRD uses MAPQ>=10 to filter Bowtie 2-aligned reads, while our pipeline only enables high-quality uniquely mapping reads for downstream analysis. We downloaded all human/mouse GTRD peak clusters called by MACS2. The peak summits were then extracted to investigate the overlap between TEs and TF bound regions (Supplementary Figure 1). For this analysis, we only kept TFs with peak clusters of more than 10,000, which resulted in 435 human and 266 mouse TFs. The fraction of TF peak summits that overlapped with TEs was then calculated for each TFs. Note that later in Figure 6, to capture all possible functional RELA-bound MER81 elements in human, we again used the RELA peak clusters documented in GTRD to expand RELA-bound MER81 instances in HAEC (n=87) to all the reported ones (n=327) for functional assessments (e.g., Figure 6B, 6C and 6D).

### TF Motif analysis

In this study, we have performed two types of motif analysis: (1) motif enrichment analysis for a group of sequences; (2) motif scan analysis for individual TE instance or consensus sequence. The vertebrate TF motifs documented in JASPAR CORE (746 TFs in total; (Castro-Mondragon et al. 2022)) were selected as the database for both analyses. (1) For motif enrichment analysis, CentriMo from MEME suite (Bailey et al. 2015) (version 4.12.0) was used with the threshold “-ethresh” set to 1. The query sequences, for example, from Figure 5, were from RELA-bound TE instances. The overlapped RELA peak summits were first extracted for all these instances and extended with 250 bp in both directions. CentriMo was then performed for instances of each TE subfamily that was identified to be significant as shown in Figure 2. From the enrichment results, top 10 enriched TFs based on E values were saved for each TE subfamily and only TEs with significant TF motifs were considered for the following analysis. To make the enrichment results comparable across different TE subfamilies, we chose to use the rank of TF motifs in each TE subfamily as the metric and visualized these values in heatmaps as shown in Figure 5A, 5B and Supplementary Figure 6. (2) In terms of motif scan analysis, FIMO from the MEME suite (version 4.12.0) was used with default parameters <u>(Grant</u> <u>et al. 2011)</u>. For example, FIMO was used to determine RELA motifs in each TE consensus sequence (Figure 2B) and MER81 instances from all the reconstructed ancestral genomes (Figure 7D). In addition, one major downstream analysis from motif scan results was the visualization of motif distribution relative to the consensus sequence (e.g., Figure 5C, 6B and 7D). RepeatMasker annotation was used to determine the corresponding position in the consensus sequence for each identified motif from instances. The occurrences of motifs in each position were then counted and normalized by the total number of instances.

### Motif-word analysis for several example TE subfamilies

Motif-word analysis has been performed for Bov-tA1/2/3 and MER41B. According to the motif distribution plots (e.g., Figure 5C and Supplementary Figure 8A), the position of the most prominent “suboptimal” RELA motif (a sequence that is only one or two nucleotides away from a matched RELA motif) in the consensus sequence was determined for each TE subfamily. Next, the RELA motif was scanned over all the instances and only matches with RELA motifs in instances that corresponded to the position of the “suboptimal” RELA motif in consensus were recorded. The recorded RELA motif-words were then counted for occurrences and the most abundant motif-words were picked for closer investigation based on the elbow plot (Supplementary Figure 7B). Finally, the occurrences of these motif-words in the queried position were counted for all the instances (denoted as “background”) and RELA-bound instances (Figure 5D and Supplementary Figure 8B).

### Visualization of sequence constraints and epigenetic signal profile

We used phastCons scores (Siepel et al. 2005) (downloaded from <u>hg38.phastCons100way.bw</u>) to evaluate the sequence constraints around MER81 instances (Figure 6C). To achieve the compatibility with phastCons scores from hg38, we lifted sequences from hg19 to hg38 using liftOver (with all default parameters) (Hinrichs et al. 2006). For epigenetic data, we used the UCSC Genome Browser (Tyner et al. 2017) to visualize the signal from ChIP-seq, ATAC-seq and loops identified from H3K4me3 HiChIP on the whole genome scale. As to the aggregate plot of ChIP-seq and ATAC-seq signal around RELA-bound TEs (e.g., Figure 3A, 6D and Supplementary Figure 4), we used computeMatrix and plotProfile functions from deepTools2 suite (Ramírez et al. 2016) (v3.1.3). To visualize stranded eRNA signals from ChRO-seq, we used HOMER (Heinz et al. 2010) (v4.11). Details and parameter choices were as Alizada et al. 2021.

### Target gene association and functional enrichment analysis for RELA-bound TEs

Human RNA-seq data have been used to determine the significantly regulated (up-regulated and down-regulated) and constitutive genes. The three gene sets used in this study were previously defined in Alizada *et al*. 2021 using both intronic and exonic reads. The distance of the four groups of RELA-bound regions relative to the three gene groups (Figure 3C) was determined using the TSS annotation of GENCODE (v19). Note that we considered merely one TSS per gene. The occurrences were counted in 10 kb bins and then converted to the corresponding fraction value for the distribution in Figure 3C. For the background reference, all the annotated TE instances were considered and the corresponding fraction was calculated accordingly. As to the functional enrichment analysis, GREAT (McLean et al. 2010) (v3.0.0) was performed for the genomic coordinates of regions of interest. For Figure 3D, to make the enrichment results comparable for the four groups of regions, a number-matched strategy was used. In brief, we selected the top 1492 RELA peaks from the conserved and human-specific groups based on peak intensity to match the total number of old TE-derived RELA bound regions (n=1492) as input for GREAT. Ontology terms from Biological Process (BP) and MSigDB pathway were extracted for each peak category. The top enriched ontology terms shown in Figure 3D were generated by combining the top 5 enriched terms (from BP/ MSigDB pathway; ranked by binomial FDR) of the four peak categories.

### Cross-species and cross-cell-type comparison of RELA-bound regions

We define conserved RELA binding events in one species as the ones that can be detected in the orthologous regions of at least another species with at least 1bp overlap. The workflow to identify the conservation degree (either conserved or species-specific) of any given RELA-bound region is as Alizada et al. 2021. Based on this rule, human RELA peaks were divided into conserved and human-specific groups, as shown in Figure 3. Similarly, when compared across human cell types, a RELA-bound region is shared if another site is identified from at least one other cell type with at least 1 bp overlap.

### Ancestral genome reconstruction, MER81 annotation and RELA motif detection

The genomes were reconstructed using Ancestors 1.1 (Diallo et al. 2010) which was applied to an alignment of 58 eutherian genomes extracted from UCSC’s 100-way whole genome alignment. Briefly, the tool employs a maximum likelihood tree-HMM approach with a context-dependent substitution algorithm to infer indels and substitutions within each branch of the phylogenetic tree and derive the ancestral sequence at each node. Its accuracy has been extensively demonstrated, especially for primate ancestors (Blanchette et al. 2004).

RepeatMasker (Tempel 2012) with Dfam libraries (Hubley et al. 2016) was run for each reconstructed genome to annotate all human transposable elements. More details were extensively described by Campitelli et al. 2022 (Campitelli et al. 2022). To acquire high-quality MER81 annotations, we additionally required MER81 hits to be marked as non-redundant and contain a minimal length of 90 bp with RepeatMasker. For the detection of RELA motifs, we extracted sequences of all the valid MER81 instances from these reconstructed genomes and used FIMO to scan RELA motifs (as described in the TF motif analysis section). The motif occurrences within identified MER81 instances for each genome were counted and visualized in Figure 7B and 7C. Furthermore, the corresponding position of motif matches relative to the consensus sequence was also recorded for each genome (Figure 7D).

### Simulation for neutral decay of MER81 elements during evolution

MER81 elements were extracted from the oldest reconstructed genome (the common ancestor of human and star-nosed mole, or node 10). A primate-specific estimate of nucleotide substitution rate (2.2 ×e-9/ (year×site); (Bulmer et al. 1991; Pace et al. 2008)) was used to simulate sequence change for the following 10 evolutionary time points. Starting from the collection of the oldest MER81 elements (real observations), all the following time points were estimated solely on nucleotide substitution. For example, each nucleotide of a MER81 instance was calculated for the likelihood of substitution based on the rate and evolutionary time, and the value of the nucleotide in the following time point/node was chosen based on the likelihood. For each node/ time point, all the simulated MER81 elements were scanned for RELA motifs using FIMO and the percentage of elements with significant motifs was recorded. We have iterated 1,000 times to obtain the mean values for each node, which were used to plot the gray line in Figure 7C.

In this study, we used several published datasets: the human/mouse/cow ChIP-seq and ATAC-seq data were downloaded from ArrayExpress with accession numbers E-MTAB-7889 and E-MTAB-7878, respectively (Alizada et al. 2021). HAEC RNA-seq and TeloHAEC ChRO-seq were downloaded from ArrayExpress with accession numbers E-MTAB-7896 and E-MTAB-9425 (Alizada et al. 2021). The RELA ChIP-seq data for HUVEC (Brown et al. 2014), LCL (Kasowski et al. 2010) and Adipocytes (Schmidt et al. 2015) were downloaded from GEO database under accession numbers: GSE54000, GSE19486 and GSE64233 respectively.

## DATA ACCESS

All raw and processed sequencing data generated in this study have been submitted to the ArrayExpress database (https://www.ebi.ac.uk/biostudies/arrayexpress/) under accession numbers E-MTAB-12212 and E-MTAB-12213. Custom scripts and files used for the analysis are provided at doi: 10.5281/zenodo.8156730 (url: https://doi.org/10.5281/zenodo.8156730)

## COMPETING INTEREST STATEMENT

The authors declare that they have no competing interests.

## ACKNOWLEDGMENTS

We would like to thank Alan Moses, Dustin Sokolowski, Cadia Chan and Huayun Hou for helpful discussions and the Centre for Applied Genomics (TCAG) for assistance with massively parallel sequencing sequencing. Funding for this project was provided by the Canadian Institutes of Health Research (CIHR) (201603PJT-364832) to M.D.W. and J.E.F. and the National Institutes of Health (S.R., M.D.W and J.A.M; R01-HG010045-01). J.E.F. and M.D.W. were supported by an Early Researcher Awards from the Ontario Ministry of Research and Innovation and Tier 2 Canada Research Chairs from CIHR. N.K. received a Canada Graduate Scholarship from the Natural Sciences and Engineering Research Council of Canada (NSERC) and an Ontario Graduate Scholarship. L.A. was supported by a NSERC CGS-M. K.R. was partially supported by a postdoctoral fellowship from the Ted Rogers Centre for Heart Research. L.C. was supported by a NSERC PGS-D scholarship.

## AUTHOR CONTRIBUTIONS

L.W. performed data analysis, wrote the manuscript. A.A. R.K. L.A. designed and performed experiments; N.K. performed luciferase assays; T.T., I.C.T. performed *CRISPR-Cas9* experiments; L.F.C., Z.P., S.P. performed ancestral genome analyses; T.H., S.R. J.M. and J.F. provided funding and supervision. M.D.W. designed experiments, wrote the manuscript, acquired funding, and supervised the project. All authors edited and approved the manuscript.

## CODE AVAILABILITY

Source code for main figures and supplementary figures is available on Wilson lab GitHub repository (https://github.com/wilsonlabgroup/RELA_TE).

## Notes

https://www.ebi.ac.uk/biostudies/arrayexpress/studies/E-MTAB-12212

https://www.ebi.ac.uk/biostudies/arrayexpress/studies/E-MTAB-12213

https://doi.org/10.5281/zenodo.8156730

### Competing Interest Statement

The authors have declared no competing interest.

### Summary of Updates

Several new analyses were performed, more information regarding data processing was given, additional CRISPR/Cas9 validation work was performed. We added additional controls to our ancestral reconstruction analysis. We investigated further the possibility that conserved orthologous NF-κB binding 'turned over' into TEs. The writing and referencing the literature has also been improved.

## REFERENCES

Alizada A, Khyzha N, Wang L, Antounians L, Chen X, Khor M, Liang M, Rathnakumar K, Weirauch MT, Medina-Rivera A, et al. 2021. Conserved regulatory logic at accessible and inaccessible chromatin during the acute inflammatory response in mammals. Nat Commun 12: 567.

Antonaki A, Demetriades C, Polyzos A, Banos A, Vatsellas G, Lavigne MD, Apostolou E, Mantouvalou E, Papadopoulou D, Mosialos G, et al. 2011. Genomic analysis reveals a novel nuclear factor-κB (NF-κB)-binding site in Alu-repetitive elements. J Biol Chem 286: 38768– 38782.

Apostolou E, Thanos D. 2008. Virus Infection Induces NF-kappaB-dependent interchromosomal associations mediating monoallelic IFN-beta gene expression. Cell 134: 85–96.

Bailey TL, Johnson J, Grant CE, Noble WS. 2015. The MEME Suite. Nucleic Acids Res 43: W39–49.

Bao W, Kojima KK, Kohany O. 2015. Repbase Update, a database of repetitive elements in eukaryotic genomes. Mob DNA 6: 11.

Bennett EA, Keller H, Mills RE, Schmidt S, Moran JV, Weichenrieder O, Devine SE. 2008. Active Alu retrotransposons in the human genome. Genome Res 18: 1875–1883.

Bhatt D, Ghosh S. 2014. Regulation of the NF-κB-Mediated Transcription of Inflammatory Genes. Front Immunol 5: 71.

Blanchette M, Green ED, Miller W, Haussler D. 2004. Reconstructing large regions of an ancestral mammalian genome in silico. Genome Res 14: 2412–2423.

Bogdan L, Barreiro L, Bourque G. 2020. Transposable elements have contributed human regulatory regions that are activated upon bacterial infection. Philos Trans R Soc Lond B Biol Sci 375: 20190332.

Bourque G, Leong B, Vega VB, Chen X, Lee YL, Srinivasan KG, Chew J-L, Ruan Y, Wei C-L, Ng HH, et al. 2008. Evolution of the mammalian transcription factor binding repertoire via transposable elements. Genome Res 18: 1752–1762.

Brown JD, Lin CY, Duan Q, Griffin G, Federation A, Paranal RM, Bair S, Newton G, Lichtman A, Kung A, et al. 2014. NF-κB directs dynamic super enhancer formation in inflammation and atherogenesis. Mol Cell 56: 219–231.

Buecker C, Wysocka J. 2012. Enhancers as information integration hubs in development: lessons from genomics. Trends Genet 28: 276–284.

Bulmer M, Wolfe KH, Sharp PM. 1991. Synonymous nucleotide substitution rates in mammalian genes: implications for the molecular clock and the relationship of mammalian orders. Proc Natl Acad Sci USA 88: 5974–5978.

Campitelli LF, Yellan I, Albu M, Barazandeh M, Patel ZM, Blanchette M, Hughes TR. 2022. Reconstruction of full-length LINE-1 progenitors from ancestral genomes. Genetics 221.

Carey KM, Patterson G, Wheeler TJ. 2021. Transposable element subfamily annotation has a reproducibility problem. Mob DNA 12: 4.

Castro-Mondragon JA, Riudavets-Puig R, Rauluseviciute I, Lemma RB, Turchi L, Blanc-Mathieu R, Lucas J, Boddie P, Khan A, Manosalva Pérez N, et al. 2022. JASPAR 2022: the 9th release of the open-access database of transcription factor binding profiles. Nucleic Acids Res 50: D165–D173.

Chen X, Pacis A, Aracena KA, Gona S, Kwan T, Groza C, Lin YL, Sindeaux R, Yotova V, Pramatarova A, et al. 2023. Transposable elements are associated with the variable response to influenza infection. Cell Genomics 3: 100292.

Chepelev I, Wei G, Wangsa D, Tang Q, Zhao K. 2012. Characterization of genome-wide enhancer-promoter interactions reveals co-expression of interacting genes and modes of higher order chromatin organization. Cell Res 22: 490–503.

Choudhary MNK, Quaid K, Xing X, Schmidt H, Wang T. 2023. Widespread contribution of transposable elements to the rewiring of mammalian 3D genomes. Nat Commun 14: 634.

Chuong EB, Elde NC, Feschotte C. 2017. Regulatory activities of transposable elements: from conflicts to benefits. Nat Rev Genet 18: 71–86.

Chuong EB, Elde NC, Feschotte C. 2016. Regulatory evolution of innate immunity through co-option of endogenous retroviruses. Science 351: 1083–1087.

Chu T, Rice EJ, Booth GT, Salamanca HH, Wang Z, Core LJ, Longo SL, Corona RJ, Chin LS, Lis JT, et al. 2018. Chromatin run-on and sequencing maps the transcriptional regulatory landscape of glioblastoma multiforme. Nat Genet 50: 1553–1564.

Correa RG, Tergaonkar V, Ng JK, Dubova I, Izpisua-Belmonte JC, Verma IM. 2004. Characterization of NF-kappa B/I kappa B proteins in zebra fish and their involvement in notochord development. Mol Cell Biol 24: 5257–5268.

Darlington GJ, Ross SE, MacDougald OA. 1998. The role of C/EBP genes in adipocyte differentiation. J Biol Chem 273: 30057–30060.

Delfino F, Walker WH. 1998. Stage-specific nuclear expression of NF-kappaB in mammalian testis. Mol Endocrinol 12: 1696–1707.

Diallo AB, Makarenkov V, Blanchette M. 2010. Ancestors 1.0: a web server for ancestral sequence reconstruction. Bioinformatics 26: 130–131.

Ding Q, Regan SN, Xia Y, Oostrom LA, Cowan CA, Musunuru K. 2013. Enhanced efficiency of human pluripotent stem cell genome editing through replacing TALENs with CRISPRs. Cell Stem Cell 12: 393–394.

Doench JG, Fusi N, Sullender M, Hegde M, Vaimberg EW, Donovan KF, Smith I, Tothova Z, Wilen C, Orchard R, et al. 2016. Optimized sgRNA design to maximize activity and minimize off-target effects of CRISPR-Cas9. Nat Biotechnol 34: 184–191.

Dorrington MG, Fraser IDC. 2019. NF-κB Signaling in Macrophages: Dynamics, Crosstalk, and Signal Integration. Front Immunol 10: 705.

Du AY, Zhuo X, Sundaram V, Jensen NO, Chaudhari HG, Saccone NL, Cohen BA, Wang T. 2022. Functional characterization of enhancer activity during a long terminal repeat’s evolution. Genome Res 32: 1840–1851.

Emera D, Wagner GP. 2012. Transformation of a transposon into a derived prolactin promoter with function during human pregnancy. Proc Natl Acad Sci USA 109: 11246–11251.

Fang R, Yu M, Li G, Chee S, Liu T, Schmitt AD, Ren B. 2016. Mapping of long-range chromatin interactions by proximity ligation-assisted ChIP-seq. Cell Res 26: 1345–1348.

Fish JE, Cantu Gutierrez M, Dang LT, Khyzha N, Chen Z, Veitch S, Cheng HS, Khor M, Antounians L, Njock M-S, et al. 2017. Dynamic regulation of VEGF-inducible genes by an ERK/ERG/p300 transcriptional network. Development 144: 2428–2444.

Freytag SO, Paielli DL, Gilbert JD. 1994. Ectopic expression of the CCAAT/enhancer-binding protein alpha promotes the adipogenic program in a variety of mouse fibroblastic cells. Genes Dev 8: 1654–1663.

Fuentes DR, Swigut T, Wysocka J. 2018. Systematic perturbation of retroviral LTRs reveals widespread long-range effects on human gene regulation. eLife 7.

Fueyo R, Judd J, Feschotte C, Wysocka J. 2022. Roles of transposable elements in the regulation of mammalian transcription. Nat Rev Mol Cell Biol 23: 481–497.

Fulco CP, Nasser J, Jones TR, Munson G, Bergman DT, Subramanian V, Grossman SR, Anyoha R, Doughty BR, Patwardhan TA, et al. 2019. Activity-by-contact model of enhancer-promoter regulation from thousands of CRISPR perturbations. Nat Genet 51: 1664–1669.

Gerondakis S, Siebenlist U. 2010. Roles of the NF-kappaB pathway in lymphocyte development and function. Cold Spring Harb Perspect Biol 2: a000182.

Heinz S, Benner C, Spann N, Bertolino E, Lin YC, Laslo P, Cheng JX, Murre C, Singh H, Glass CK. 2010. Simple combinations of lineage-determining transcription factors prime cis-regulatory elements required for macrophage and B cell identities. Mol Cell 38: 576–589.

Hermant C, Torres-Padilla M-E. 2021. TFs for TEs: the transcription factor repertoire of mammalian transposable elements. Genes Dev 35: 22–39.

Hetru C, Hoffmann JA. 2009. NF-kappaB in the immune response of Drosophila. Cold Spring Harb Perspect Biol 1: a000232.

Hinrichs AS, Karolchik D, Baertsch R, Barber GP, Bejerano G, Clawson H, Diekhans M, Furey TS, Harte RA, Hsu F, et al. 2006. The UCSC Genome Browser Database: update 2006. Nucleic Acids Res 34: D590–8.

Hirakawa M, Nishihara H, Kanehisa M, Okada N. 2009. Characterization and evolutionary landscape of AmnSINE1 in Amniota genomes. Gene 441: 100–110.

Hogan NT, Whalen MB, Stolze LK, Hadeli NK, Lam MT, Springstead JR, Glass CK, Romanoski CE. 2017. Transcriptional networks specifying homeostatic and inflammatory programs of gene expression in human aortic endothelial cells. eLife 6.

Hsu PD, Scott DA, Weinstein JA, Ran FA, Konermann S, Agarwala V, Li Y, Fine EJ, Wu X, Shalem O, et al. 2013. DNA targeting specificity of RNA-guided Cas9 nucleases. Nat Biotechnol 31: 827–832.

Hubley R, Finn RD, Clements J, Eddy SR, Jones TA, Bao W, Smit AFA, Wheeler TJ. 2016. The Dfam database of repetitive DNA families. Nucleic Acids Res 44: D81–9.

Ito J, Sugimoto R, Nakaoka H, Yamada S, Kimura T, Hayano T, Inoue I. 2017. Systematic identification and characterization of regulatory elements derived from human endogenous retroviruses. PLoS Genet 13: e1006883.

Jacques P-É, Jeyakani J, Bourque G. 2013. The majority of primate-specific regulatory sequences are derived from transposable elements. PLoS Genet 9: e1003504.

Judd J, Sanderson H, Feschotte C. 2021. Evolution of mouse circadian enhancers from transposable elements. Genome Biol 22: 193.

Juric I, Yu M, Abnousi A, Raviram R, Fang R, Zhao Y, Zhang Y, Qiu Y, Yang Y, Li Y, et al. 2019. MAPS: Model-based analysis of long-range chromatin interactions from PLAC-seq and HiChIP experiments. PLoS Comput Biol 15: e1006982.

Kaikkonen MU, Spann NJ, Heinz S, Romanoski CE, Allison KA, Stender JD, Chun HB, Tough DF, Prinjha RK, Benner C, et al. 2013. Remodeling of the enhancer landscape during macrophage activation is coupled to enhancer transcription. Mol Cell 51: 310–325.

Kasowski M, Grubert F, Heffelfinger C, Hariharan M, Asabere A, Waszak SM, Habegger L, Rozowsky J, Shi M, Urban AE, et al. 2010. Variation in transcription factor binding among humans. Science 328: 232–235.N

Kelly CJ, Chitko-McKown CG, Chuong EB. 2022. Ruminant-specific retrotransposons shape regulatory evolution of bovine immunity. Genome Res.

Kolmykov S, Yevshin I, Kulyashov M, Sharipov R, Kondrakhin Y, Makeev VJ, Kulakovskiy IV, Kel A, Kolpakov F. 2021. GTRD: an integrated view of transcription regulation. Nucleic Acids Res 49: D104–D111.

Kolovos P, Georgomanolis T, Koeferle A, Larkin JD, Brant L, Nikolicć M, Gusmao EG, Zirkel A, Knoch TA, van Ijcken WF, et al. 2016. Binding of nuclear factor κB to noncanonical consensus sites reveals its multimodal role during the early inflammatory response. Genome Res 26: 1478–1489.

Kumar S, Stecher G, Suleski M, Hedges SB. 2017. Timetree: A resource for timelines, timetrees, and divergence times. Mol Biol Evol 34: 1812–1819.

Kunarso G, Chia N-Y, Jeyakani J, Hwang C, Lu X, Chan Y-S, Ng H-H, Bourque G. 2010. Transposable elements have rewired the core regulatory network of human embryonic stem cells. Nat Genet 42: 631–634.

Lee HJ, Hou Y, Maeng JH, Shah NM, Chen Y, Lawson HA, Yang H, Yue F, Wang T. 2022. Epigenomic analysis reveals prevalent contribution of transposable elements to cis-regulatory elements, tissue-specific expression, and alternative promoters in zebrafish. Genome Res 32: 1424–1436.

Lilienbaum A, Sage J, Mémet S, Rassoulzadegan M, Cuzin F, Israël A. 2000. NF-κB is developmentally regulated during spermatogenesis in mice. Developmental Dynamics.

Liu T, Zhang L, Joo D, Sun S-C. 2017. NF-κB signaling in inflammation. Signal Transduct Target Ther 2: 17023.

Lynch VJ, Leclerc RD, May G, Wagner GP. 2011. Transposon-mediated rewiring of gene regulatory networks contributed to the evolution of pregnancy in mammals. Nat Genet 43: 1154–1159.

MacArthur S, Brookfield JFY. 2004. Expected rates and modes of evolution of enhancer sequences. Mol Biol Evol 21: 1064–1073.

Macfarlan TS, Gifford WD, Agarwal S, Driscoll S, Lettieri K, Wang J, Andrews SE, Franco L, Rosenfeld MG, Ren B, et al. 2011. Endogenous retroviruses and neighboring genes are coordinately repressed by LSD1/KDM1A. Genes Dev 25: 594–607.

Mali P, Yang L, Esvelt KM, Aach J, Guell M, DiCarlo JE, Norville JE, Church GM. 2013. RNA-guided human genome engineering via Cas9. Science 339: 823–826.

McLean CY, Bristor D, Hiller M, Clarke SL, Schaar BT, Lowe CB, Wenger AM, Bejerano G. 2010. GREAT improves functional interpretation of cis-regulatory regions. Nat Biotechnol 28: 495–501.

Michida H, Imoto H, Shinohara H, Yumoto N, Seki M, Umeda M, Hayashi T, Nikaido I, Kasukawa T, Suzuki Y, et al. 2020. The Number of Transcription Factors at an Enhancer Determines Switch-like Gene Expression. Cell Rep 31: 107724.

Mifsud B, Tavares-Cadete F, Young AN, Sugar R, Schoenfelder S, Ferreira L, Wingett SW, Andrews S, Grey W, Ewels PA, et al. 2015. Mapping long-range promoter contacts in human cells with high-resolution capture Hi-C. Nat Genet 47: 598–606.

Mumbach MR, Rubin AJ, Flynn RA, Dai C, Khavari PA, Greenleaf WJ, Chang HY. 2016. HiChIP: efficient and sensitive analysis of protein-directed genome architecture. Nat Methods 13: 919–922.

Muñoz-López M, García-Pérez JL. 2010. DNA transposons: nature and applications in genomics. Curr Genomics 11: 115–128.

Natoli G, Saccani S, Bosisio D, Marazzi I. 2005. Interactions of NF-kappaB with chromatin: the art of being at the right place at the right time. Nat Immunol 6: 439–445.

Nijman IJ, van Tessel P, Lenstra JA. 2002. SINE retrotransposition during the evolution of the Pecoran ruminants. J Mol Evol 54: 9–16.

Nilsson MA, Klassert D, Bertelsen MF, Hallström BM, Janke A. 2012. Activity of ancient RTE retroposons during the evolution of cows, spiral-horned antelopes, and Nilgais (Bovinae). Mol Biol Evol 29: 2885–2888.

Nishihara H, Smit AFA, Okada N. 2006. Functional noncoding sequences derived from SINEs in the mammalian genome. Genome Res 16: 864–874.

Notwell JH, Chung T, Heavner W, Bejerano G. 2015. A family of transposable elements co-opted into developmental enhancers in the mouse neocortex. Nat Commun 6: 6644.

Oeckinghaus A, Hayden MS, Ghosh S. 2011. Crosstalk in NF-κB signaling pathways. Nat Immunol 12: 695–708.

Ostuni R, Piccolo V, Barozzi I, Polletti S, Termanini A, Bonifacio S, Curina A, Prosperini E, Ghisletti S, Natoli G. 2013. Latent enhancers activated by stimulation in differentiated cells. Cell 152: 157–171.

Pace JK, Gilbert C, Clark MS, Feschotte C. 2008. Repeated horizontal transfer of a DNA transposon in mammals and other tetrapods. Proc Natl Acad Sci USA 105: 17023–17028.

Paciolla M, Boni R, Fusco F, Pescatore A, Poeta L, Ursini MV, Lioi MB, Miano MG. 2011. Nuclear factor-kappa-B-inhibitor alpha (NFKBIA) is a developmental marker of NF-κB/p65 activation during in vitro oocyte maturation and early embryogenesis. Hum Reprod 26: 1191– 1201.

Pereira JP, Kelly LM, Cyster JG. 2010. Finding the right niche: B-cell migration in the early phases of T-dependent antibody responses. Int Immunol 22: 413–419.

Pontis J, Planet E, Offner S, Turelli P, Duc J, Coudray A, Theunissen TW, Jaenisch R, Trono D. 2019. Hominoid-Specific Transposable Elements and KZFPs Facilitate Human Embryonic Genome Activation and Control Transcription in Naive Human ESCs. Cell Stem Cell 24: 724–735.e5.

Ramírez F, Ryan DP, Grüning B, Bhardwaj V, Kilpert F, Richter AS, Heyne S, Dündar F, Manke T. 2016. deepTools2: a next generation web server for deep-sequencing data analysis. Nucleic Acids Res 44: W160–5.

Rasoulpour RJ, Boekelheide K. 2005. NF-kappaB is activated in the rat testis following exposure to mono-(2-ethylhexyl) phthalate. Biol Reprod 72: 479–486.

Romanish MT, Lock WM, van de Lagemaat LN, Dunn CA, Mager DL. 2007. Repeated recruitment of LTR retrotransposons as promoters by the anti-apoptotic locus NAIP during mammalian evolution. PLoS Genet 3: e10.

Sasaki T, Nishihara H, Hirakawa M, Fujimura K, Tanaka M, Kokubo N, Kimura-Yoshida C, Matsuo I, Sumiyama K, Saitou N, et al. 2008. Possible involvement of SINEs in mammalian-specific brain formation. Proc Natl Acad Sci USA 105: 4220–4225.

Satriano J, Schlondorff D. 1994. Activation and attenuation of transcription factor NF-kB in mouse glomerular mesangial cells in response to tumor necrosis factor-alpha, immunoglobulin G, and adenosine 3’:5’-cyclic monophosphate. Evidence for involvement of reactive oxygen species. J Clin Invest 94: 1629–1636.

Schmidt D, Schwalie PC, Wilson MD, Ballester B, Gonçalves A, Kutter C, Brown GD, Marshall A, Flicek P, Odom DT. 2012. Waves of retrotransposon expansion remodel genome organization and CTCF binding in multiple mammalian lineages. Cell 148: 335–348.

Schmidt D, Wilson MD, Spyrou C, Brown GD, Hadfield J, Odom DT. 2009. ChIP-seq: using high-throughput sequencing to discover protein-DNA interactions. Methods 48: 240–248.

Schmidt SF, Larsen BD, Loft A, Nielsen R, Madsen JGS, Mandrup S. 2015. Acute TNF-induced repression of cell identity genes is mediated by NFκB-directed redistribution of cofactors from super-enhancers. Genome Res 25: 1281–1294.

Senft AD, Macfarlan TS. 2021. Transposable elements shape the evolution of mammalian development. Nat Rev Genet 22: 691–711.

Sen R, Baltimore D. 1986. Inducibility of kappa immunoglobulin enhancer-binding protein Nf-kappa B by a posttranslational mechanism. Cell 47: 921–928.

Sexton CE, Han MV. 2019. Paired-end mappability of transposable elements in the human genome. Mob DNA 10: 29.

Shah AV, Birdsey GM, Randi AM. 2016. Regulation of endothelial homeostasis, vascular development and angiogenesis by the transcription factor ERG. Vascul Pharmacol 86: 3–13.

Siepel A, Bejerano G, Pedersen JS, Hinrichs AS, Hou M, Rosenbloom K, Clawson H, Spieth J, Hillier LW, Richards S, et al. 2005. Evolutionarily conserved elements in vertebrate, insect, worm, and yeast genomes. Genome Res 15: 1034–1050.

Sundaram V, Cheng Y, Ma Z, Li D, Xing X, Edge P, Snyder MP, Wang T. 2014. Widespread contribution of transposable elements to the innovation of gene regulatory networks. Genome Res 24: 1963–1976.

Sundaram V, Wysocka J. 2020. Transposable elements as a potent source of diverse cis-regulatory sequences in mammalian genomes. Philos Trans R Soc Lond B Biol Sci 375: 20190347.

Tabruyn SP, Griffioen AW. 2007. A new role for NF-kappaB in angiogenesis inhibition. Cell Death Differ 14: 1393–1397.

Tempel S. 2012. Using and understanding RepeatMasker. Methods Mol Biol 859: 29–51.

Todd CD, Deniz Ö, Taylor D, Branco MR. 2019. Functional evaluation of transposable elements as enhancers in mouse embryonic and trophoblast stem cells. eLife 8.

Trizzino M, Park Y, Holsbach-Beltrame M, Aracena K, Mika K, Caliskan M, Perry GH, Lynch VJ, Brown CD. 2017. Transposable elements are the primary source of novelty in primate gene regulation. Genome Res 27: 1623–1633.

Tyner C, Barber GP, Casper J, Clawson H, Diekhans M, Eisenhart C, Fischer CM, Gibson D, Gonzalez JN, Guruvadoo L, et al. 2017. The UCSC Genome Browser database: 2017 update. Nucleic Acids Res 45: D626–D634.

Ward MC, Wilson MD, Barbosa-Morais NL, Schmidt D, Stark R, Pan Q, Schwalie PC, Menon S, Lukk M, Watt S, et al. 2013. Latent regulatory potential of human-specific repetitive elements. Mol Cell 49: 262–272.

Wells JN, Feschotte C. 2020. A field guide to eukaryotic transposable elements. Annu Rev Genet 54: 539–561.

Xie D, Chen C-C, Ptaszek LM, Xiao S, Cao X, Fang F, Ng HH, Lewin HA, Cowan C, Zhong S. 2010. Rewirable gene regulatory networks in the preimplantation embryonic development of three mammalian species. Genome Res 20: 804–815.

Zemojtel T, Kielbasa SM, Arndt PF, Behrens S, Bourque G, Vingron M. 2011. CpG deamination creates transcription factor-binding sites with high efficiency. Genome Biol Evol 3: 1304– 1311.

Zhang Q, Lenardo MJ, Baltimore D. 2017. 30 Years of NF-κB: A Blossoming of Relevance to Human Pathobiology. Cell 168: 37–57.

Zhang Y, Liu T, Meyer CA, Eeckhoute J, Johnson DS, Bernstein BE, Nusbaum C, Myers RM, Brown M, Li W, et al. 2008. Model-based analysis of ChIP-Seq (MACS). Genome Biol 9: R137.

