## Supplementary Figures for "Multi-species analysis of inflammatory response elements reveals ancient and lineage-specific contributions of transposable elements to NF-κB binding"

**Supplemental Figures and Tables.**

**Table of Contents:**

Supplemental Figure 1 - Transposable elements largely contribute to transcription factor bound regions in the three species.

Supplemental Figure 2 - TEs primarily derive species-specific RELA-bound regions and potentially contribute to micro-turnover of binding events.

Supplemental Figure 3 - Heatmaps of several epigenetic features around TE-derived RELA-bound regions.

Supplemental Figure 4 - TE-derived RELA-bound regions exhibit active enhancer features in aortic endothelial cells from mouse and cow.

Supplemental Figure 5 - Conservation degree vs cross cell type usage of RELA-bound regions derived from significantly enriched TEs in human.

Supplemental Figure 6 - Heatmaps of significantly enriched TF motifs.

Supplemental Figure 7 - Motif-word analysis for selected transposons.

Supplemental Figure 8 - Epigenetic profile and functional enrichment of MER81-derived RELA-bound regions in human aortic endothelial cells.

Supplemental Figure 9 - Luciferase experiments and CRISPR-Cas9 deletion experiments of MER81 elements.

Supplemental Figure 10 - Ancestral reconstruction of RELA/JUN motif changes within significantly enriched human TE subfamilies.

Supplemental Table 1 - MER81 annotations by RepeatMasker in 24 selected mammals.

Supplemental Table 2 - Examples of RELA bound MER81 instances near interferon signaling related genes.

Supplemental Table 3 - Primer sequences and gRNAs.

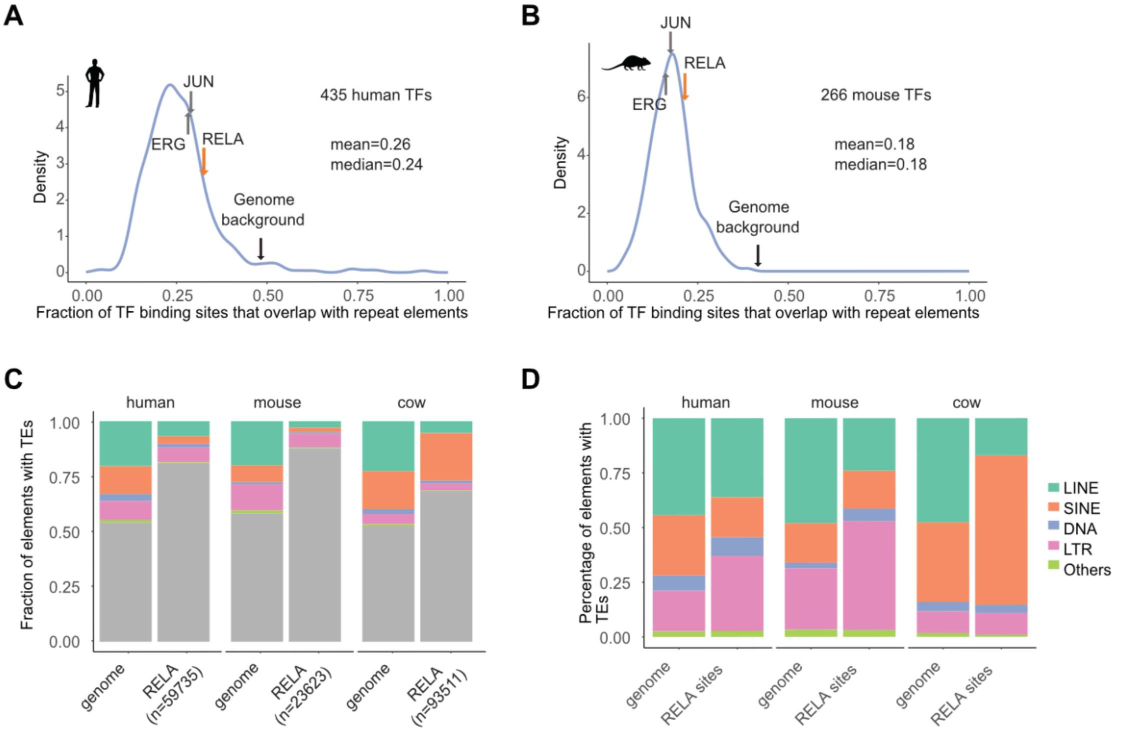


**Supplemental Figure 1**. Transposable elements largely contribute to transcription factor bound regions in the three species. Histogram of the overlap fraction of RepeatMasker annotated TEs and the peak summits of all investigated TFs from A) human and B) mouse from GTRD database (Kolmykov et al. 2021). The orange arrow indicates the overlap fraction of TEs and RELA peak summits and the black arrow indicates the expected overlap fraction based on the genome coverage of TEs. Two additional TFs (ERG and JUN) were shown as well. C) The overlap between RELA-bound regions (our RELA ChIP-seq data) and four major TE classes. For each species, RELA bound regions/peaks were identified by calling peaks on the stimulated samples using the unstimulated sample as input via MACS2. We define an overlap if a RELA peak summit intersects with a TE. Genome coverage of TE classes in each species serves as a reference. D) The composition of TEs within the whole genome and RELA-bound regions.

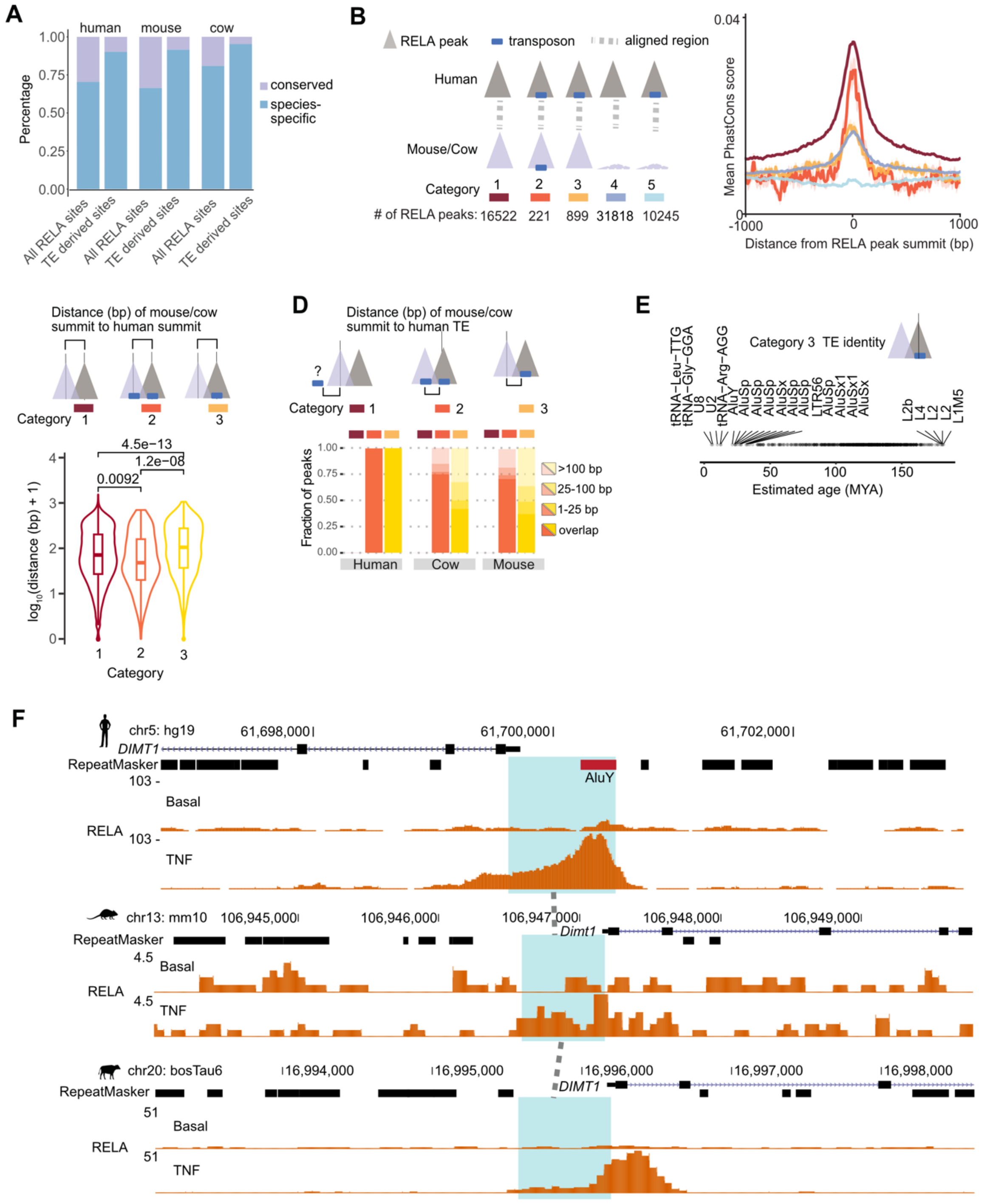
A
human
mouse
cow
Percentage
conserved
species-specific
All RELA sites
TE derived sites
B
RELA peak
transposon
aligned region
Human
Mouse/Cow
Category
### of RELA peaks: 16522 221 899 31818 10245
Mean PhastCons score
Distance from RELA peak summit (bp)
C
Distance (bp) of mouse/cow summit to human summit
Category
log10(distance (bp) + 1)
4.5e−13
1.2e−08
0.0092
D
Distance (bp) of mouse/cow summit to human TE
Fraction of peaks
>100 bp
25-100 bp
1-25 bp
overlap
Human
Cow
Mouse
E
Category 3 TE identity
tRNA−Leu−TTG
tRNA−Gly−GGA
U8
U2
tRNA−Arg−AGG
AluY
AluSp
AluSp
AluSx
AluSp
AluSp
LTR56
AluSp
AluSx1
AluSx1
AluSx
L2b
L4
L2
L1M5
Estimated age (MYA)
F
chr5: hg19
61,698,000
61,700,000
61,702,000
DIMT1
RepeatMasker
AluY
Basal
RELA
TNF
chr13: mm10
106,945,000
106,946,000
106,947,000
106,948,000
106,949,000
Dimt1
chr20: bosTau6
16,994,000
16,995,000
16,996,000
16,997,000
16,998,000

**Supplemental Figure 2**. TEs primarily derive species-specific RELA-bound regions and potentially contribute to micro-turnover of binding events. A) TEs are associated with species-specific RELA-bound regions. The conservation of a RELA-bound region is defined if another RELA-bound region can be detected in the orthologous region of another species with at least 1 bp overlap. B) Sequence constraints over a series of RELA peak categories. The top panel illustrates five different RELA peak categories that involve a TE or not; The bottom panel shows mean PhastCons scores around the five categories with the RELA peak summits as the visualization center. C) The "micro-shift" of RELA peak summits of conserved RELA binding events. RELA peak summits from the three species were lifted to the coordinates of their common ancestor sequences. Lifted summits from mouse/cow were then compared with their counterparts from human to calculate the shifted distance. Note that only the first three categories from B) were considered in the analysis. D) "Re-annotation" of the sequence identity for lifted mouse/cow RELA peak summits. Human TE annotation by RepeatMasker was utilized to annotate lifted mouse/cow RELA peak summits to examine a possibility of TE mis-annotations or potential turnover events. E) Highlighted TE instances that overlap human RELA peak summits of category 3. F) A potential candidate of TE-derived micro-turnover in human. A human-cow conserved RELA binding event locates on the promoter region of gene *DIMT1*. Note that a human-specific TE, AluY, possibly drives a "micro-turnover" of this binding event, despite the overall conservation of activity within this region. The green shade denotes orthologous sequences determined by our pipeline.

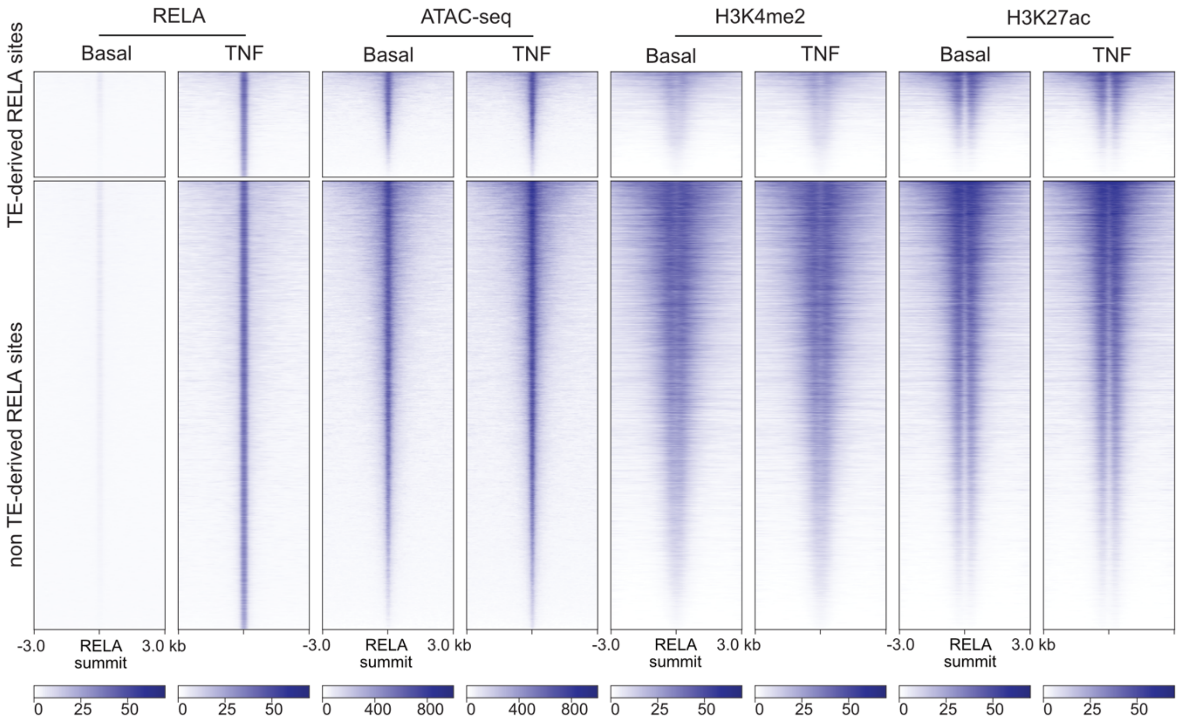


**Supplemental Figure 3**. Heatmaps of several epigenetic features around TE-derived RELA-bound regions. The heatmaps include RELA, H3K4me2, H3K27ac ChIP-seq as well as ATAC-seq from human aortic endothelial cells with/ without TNF stimulation for 45 min. The signal was plotted over TE-derived vs non TE-derived RELA-bound regions.

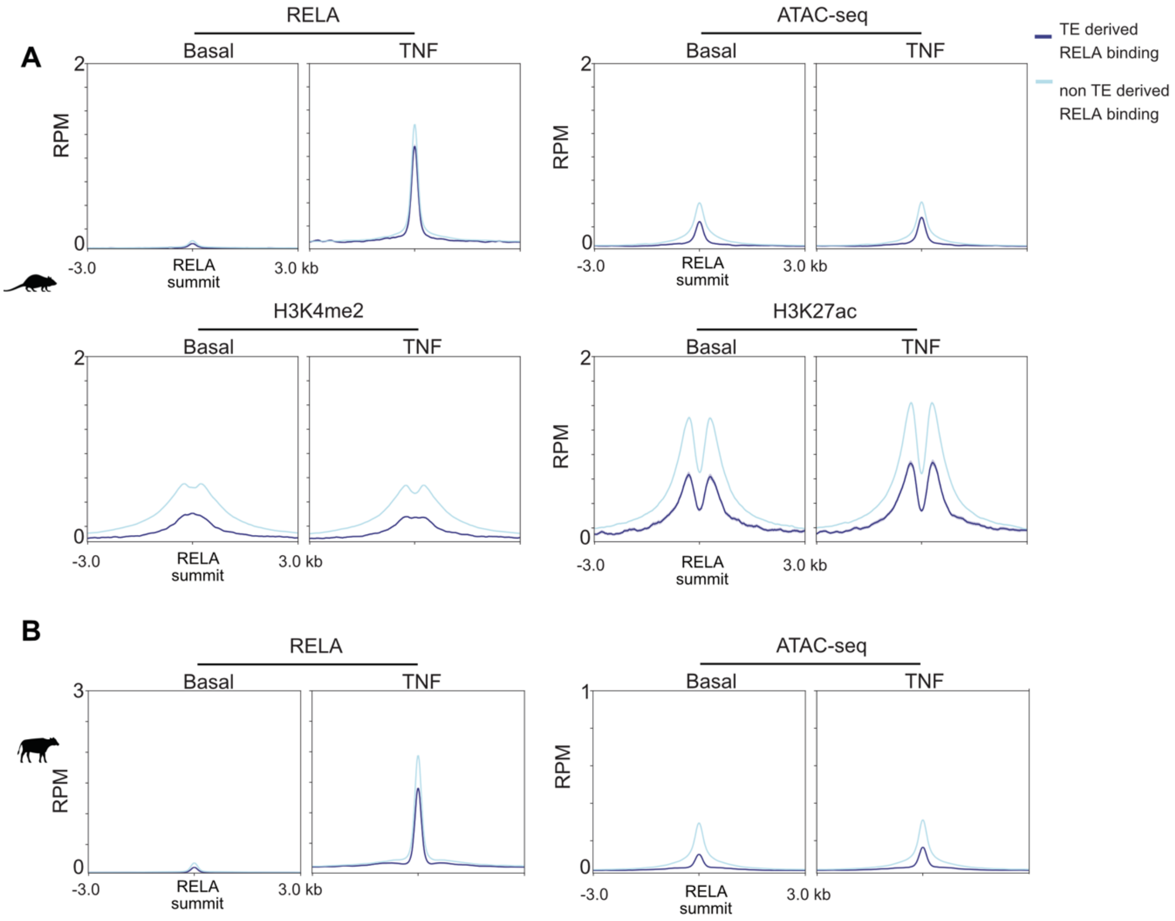


**Supplemental Figure 4**. TE-derived RELA-bound regions exhibit active enhancer features in aortic endothelial cells from mouse and cow. A) Aggregate plots of RELA, H3K4me2 and H3K27ac ChIP-seq as well as ATAC-seq around TE-derived RELA-bound regions in mouse. B) Aggregate plots of RELA ChIP-seq and ATAC-seq around TE-derived RELA-bound regions in cow.

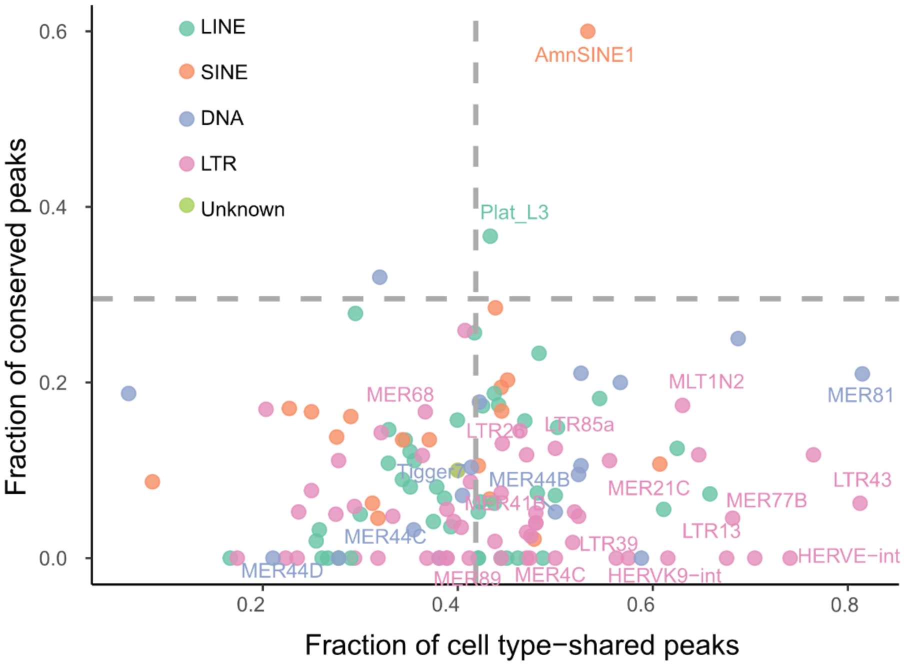


**Supplemental Figure 5**. Conservation degree vs cross cell type usage of RELA-bound regions derived from significantly enriched TEs in human. Significantly enriched TE subfamilies identified from human aortic endothelial cells are labeled. The expected fraction of conserved peaks (horizontal gray line) was calculated based on the fraction of the conserved orthologous RELA peaks from human relative to mouse and cow. The expected fraction of cell type shared peaks (vertical gray line) was obtained from human aortic endothelial cells relative to all the other three cell types.

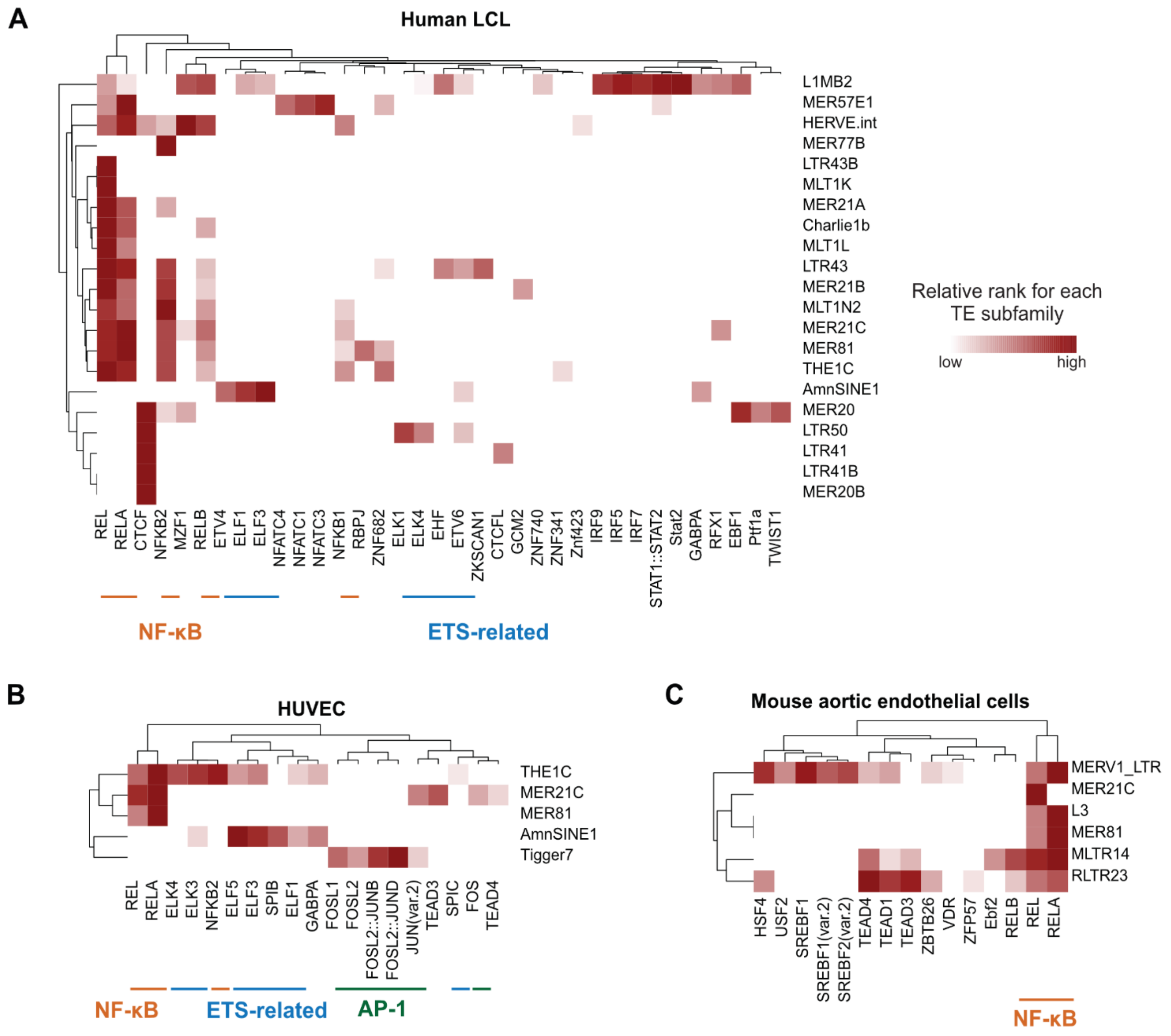


**Supplemental Figure 6.** Heatmaps of significantly enriched TF motifs. A) human LCL, B) HUVEC and C) mouse aortic endothelial cells. Significantly enriched TE subfamilies in the three cell types were investigated and only TEs detected with significant TF motifs are shown in the heatmaps. The intensity of the heatmap represents the rank of significantly enriched TF motifs for each TE (see more details in Methods). TF members of NF-κB, ETS and AP-1 have been highlighted.

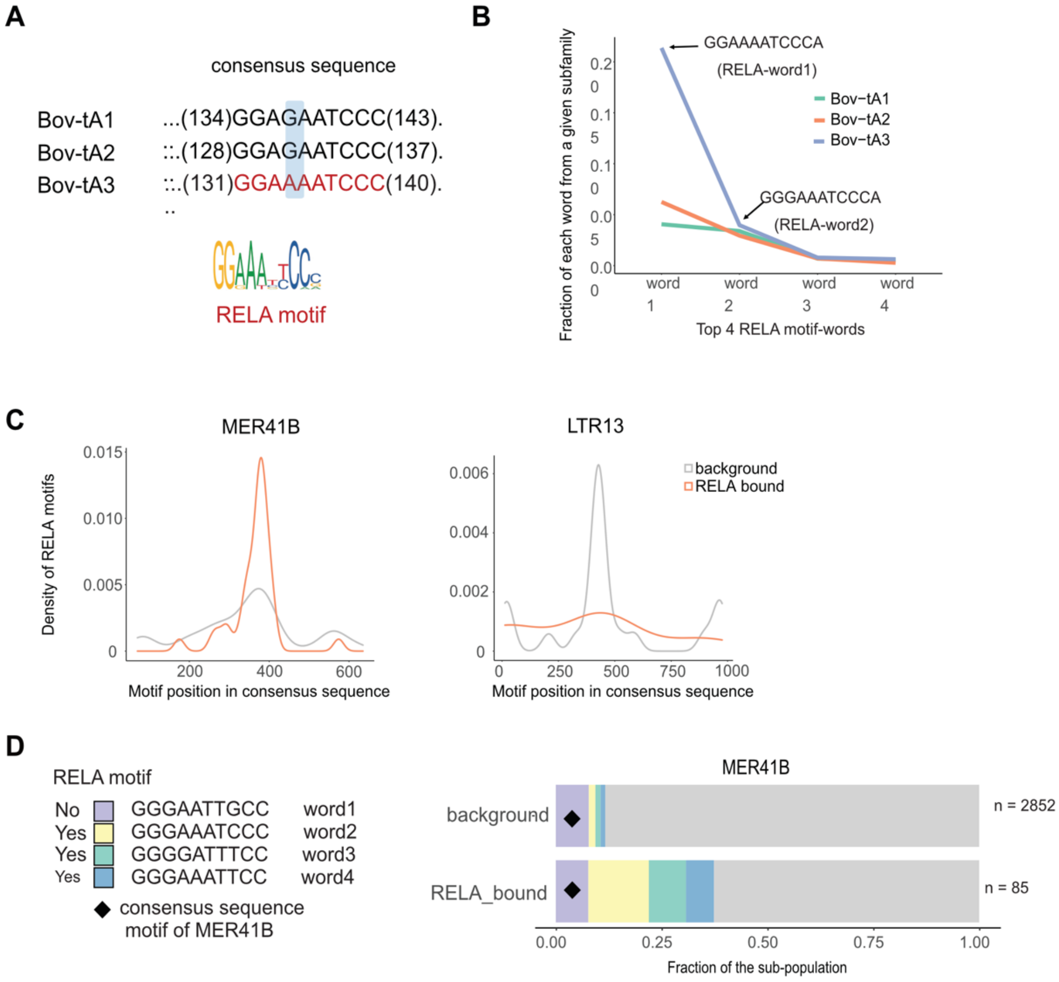


**Supplemental Figure 7**. Motif-word analysis for selected transposons: A) and B) for Bov-tA1/2/3 as well as C) and D) for MER41B. A) Part of aligned consensus sequences from Bov-tA1/2/3. Note only Bov-tA3 sequence matches with the RELA motif at this position because of a single nucleotide difference. B) The fraction of the instances that match with the top 4 most abundant RELA motif-words at the investigated position of Bov-tA1/2/3. Note the top two RELA motif-words can account for the majority of the RELA motifs at the investigated position. The exact sequences of the top two RELA motif-words are labeled. These two sequences are shared across all three subfamilies. C) RELA motif distribution relative to the consensuses of MER41B and LTR13. Gray and orange represent all instances and RELA-bound instances, respectively. D) Fine stratification of MER41B instances based on abundant RELA motif-words. The left panel shows four investigated motif-words, including a consensus word of MER41B (word1), and the top three abundant RELA motif-words (word2, word3 and word4). The right panel displays the fraction of instances that match with each word for all instances and RELA-bound instances. Grey represents all the other sequences. The number of instances of all and RELA-bound MER41B instances are labeled next to the bar.

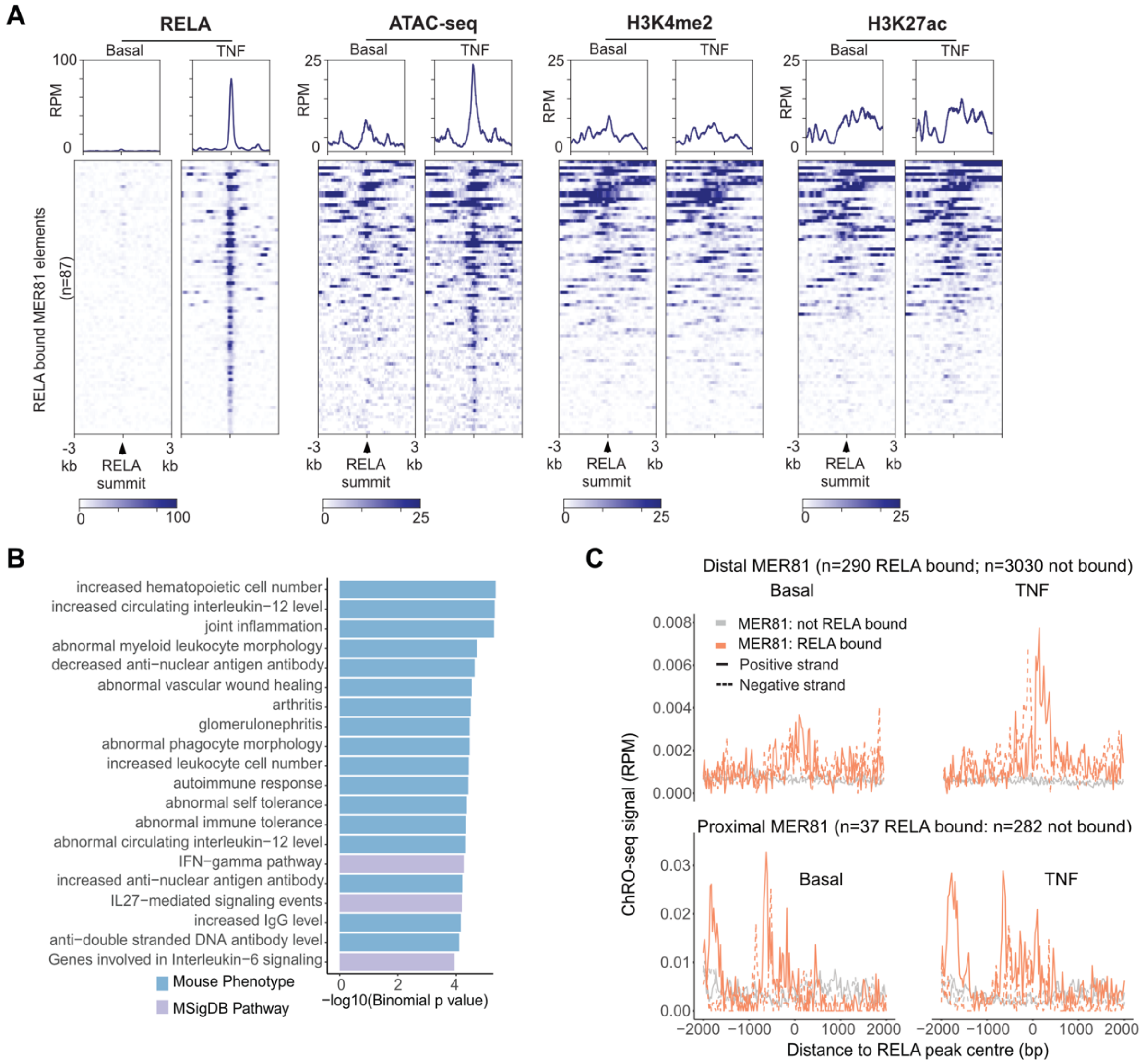


**Supplemental Figure 8**. Epigenetic profile and functional enrichment of MER81-derived RELA-bound regions in human aortic endothelial cells (n=87). A) Aggregate plots and corresponding heatmaps of RELA, H3K4me2 and H3K27ac ChIP-seq as well as ATAC-seq around MER81-derived RELA-bound regions in human aortic endothelial cells. B) Functional enrichment results from MER81-derived RELA-bound regions by GREAT. C) eRNA signals on RELA bound/non-bound MER81 instances. eRNA signals from TeloHAEC ChRO-seq data were utilized for the visualization. MER81 instances are divided into distal or proximal elements based on the distance relative to the nearest annotated gene TSS (cutoff: 3kb).

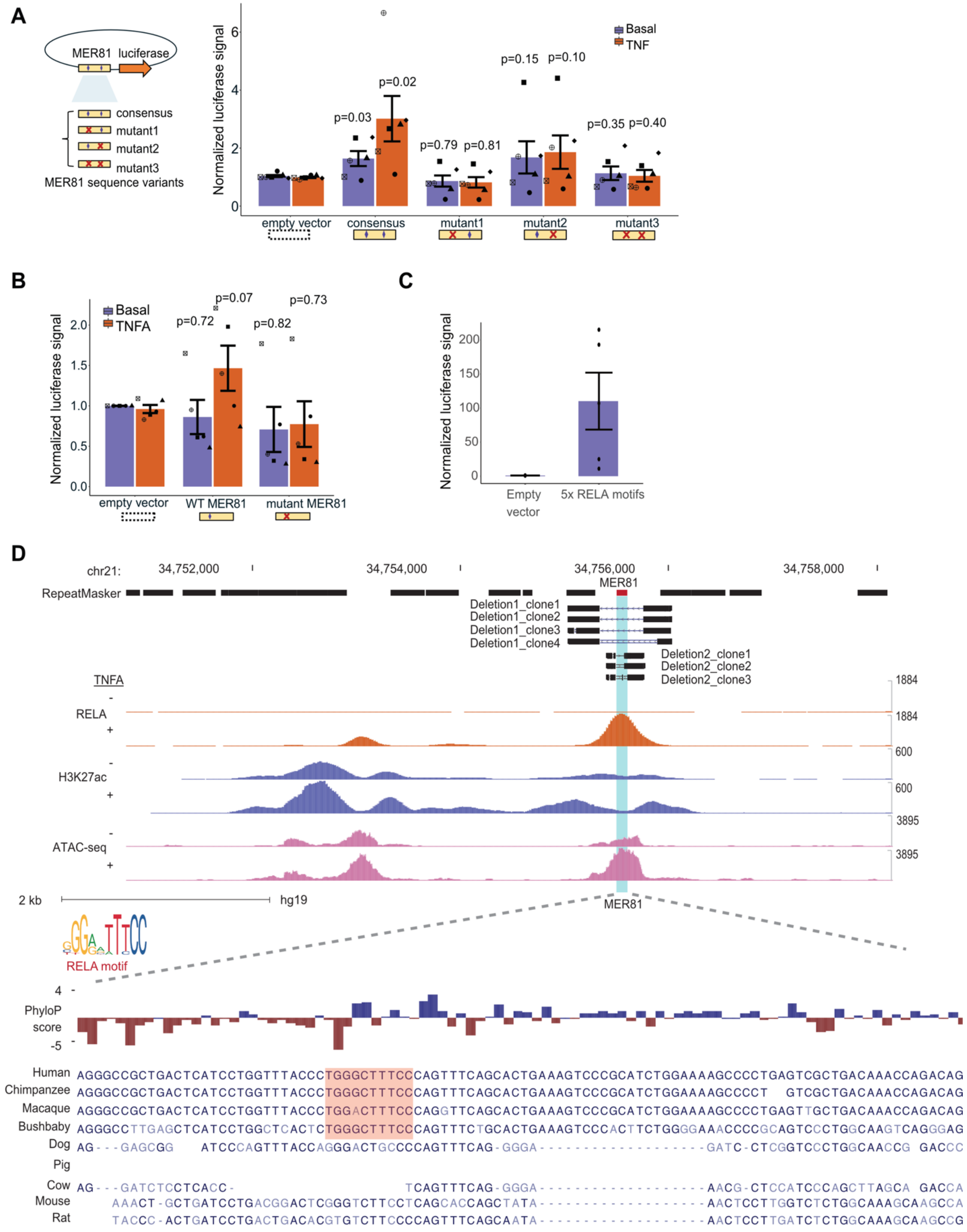


**Supplemental Figure 9**. Luciferase experiments and *CRISPR-Cas9* deletion experiments of MER81 elements. A) Functional validation of the enhancer activity of MER81 consensus sequence. Luciferase reporter assays were performed to determine the enhancer activity with/without TNF stimulation for 3 hours in HEK293T cells. The figure shows luciferase signals of MER81 consensus sequence and three sequence variants. B) The luciferase signals of the native MER81 sequence shown in Figure 6A and its variant sequence. Luciferase reporter assays were performed with/without TNF stimulation for 3 hours in HEK293T cells. C) The controls of luciferase experiments under the unstimulated condition. We originally included a positive control that contains 5 × tandem RELA motifs in the experiment. A motif scan test was performed using 746 known vertebrate TF motifs documented in JASPAR (Castro-Mondragon et al. 2022) to make sure no new TF motifs occur within the newly designed sequences. All these sequences of interest were directly synthesized and cloned in pGL3 luciferase construct backbone (service from General Biosystems). Since NF-κB signaling is pervasive across cell types, we chose a simple cell line, HEK293T, as the model to test the enhancer activity of the sequences. HEK293T cells (ATCC, cat# CRL-1573) were cultured in DMEM media (Gibco, cat# 11965092) supplemented with 10% FBS (Gibco, cat# 12483020). For luciferase assays, cells were seeded in 12-well dishes with a technical triplicate for each tested condition. HEK293Ts were transfected at 80% confluency with 1ug of pGL3 luciferase constructs and 0.1ug of pRenilla using Lipofectamine 2000 (2ul/1ug DNA, Invitrogen) per well in 1mL of optiMEM. After 5 hours, optiMEM was replaced with 2mL of DMEM + 10% FBS. For TNF treatment, 10 ng/ml recombinant human TNF (Cell Applications, cat# RP1111-50) was added for 3 hours. Luciferase reporter assay was performed using the Dual-Luciferase Reporter Assay System (Promega). 24h post-transfection, media was aspirated and 250 µl of passive lysis buffer (1X) was added to each well. Lysis was performed by shaking at 20 min covered in aluminum foil at RT. The GloMax20/20 Luminometer (Promega) was used to measure dual luciferase (*Renilla* and Firefly). Prior to taking measurements, lysed samples were diluted 10-fold to be within a detectable range of the luminometer. Afterward, 20 µl of diluted sample was added to each Eppendorf tube and samples were processed one by one. First 100 µlof the LARII reagent was added and the Firefly luciferase reading was obtained. Immediately after, 100 µl of the S&G reagent was added and the *Renilla* luciferase reading was obtained. To analyze the data, the Firefly/*Renilla* luciferase value was obtained. *Renilla* only background values were subtracted. All data points were then normalized to empty pGL3 control. D) Genome browser view of CRISPR deleted regions of the selected native MER81 instance in human. Shown are signals of RELA, H3K27ac ChIP-seq and ATAC-seq in human as well as the multiple sequence alignment from the 241-way Cactus alignment for selected species. "-" and "+" denote basal and TNF stimulation, respectively. For A, B and C, we have performed unpaired one-sided *t*-test for each group compared to the empty vector under the corresponding condition. Significance values of the statistical tests are shown on top of each bar.

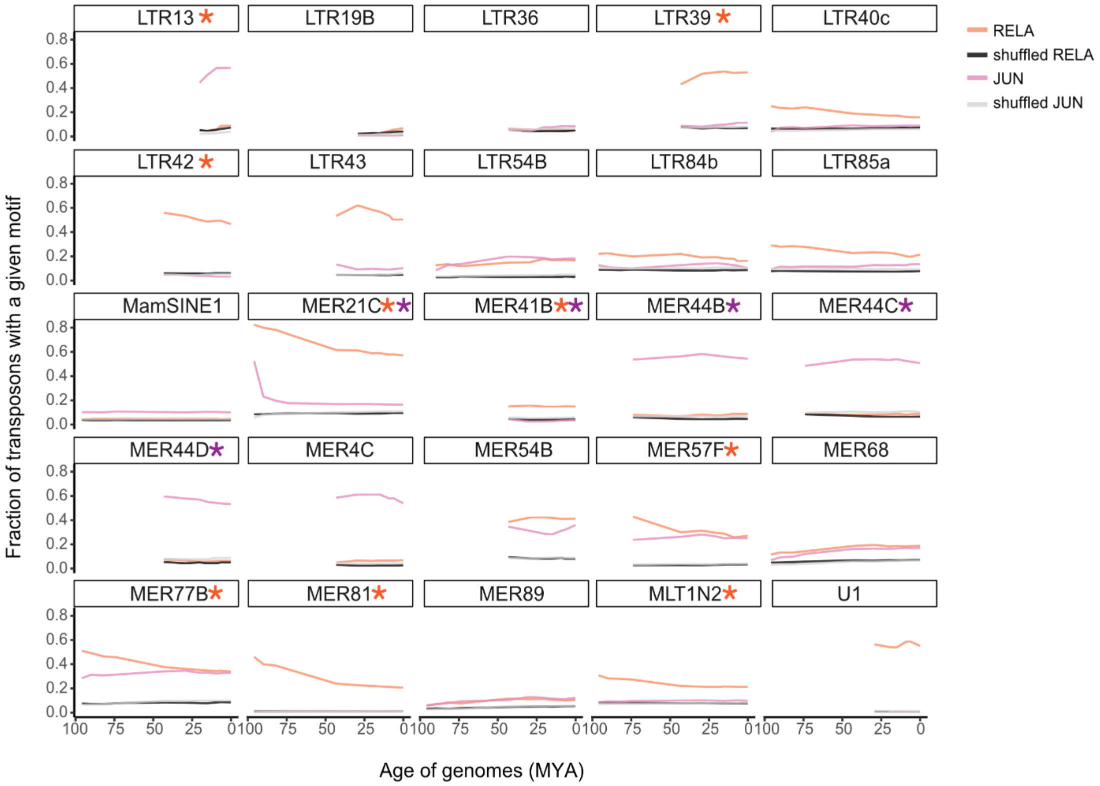


**Supplemental Figure 10**. Ancestral reconstruction of RELA/JUN motif changes within significantly enriched human TE subfamilies. The proportion of TE subfamily that contains TF motifs (RELA, JUN, or the corresponding shuffled motifs) is shown as a function of the estimated evolutionary age of each node/common ancestor from Figure 7A (divergence time estimation obtained from TIMETREE database (Kumar et al. 2017)). For shuffled motifs, 100 random shuffles (with a Pearson correlation value < 0.05 compared to the original motif) were used as a control set for RELA/JUN motifs. The median motif matches from the 100 random shuffles were used and visualized for each TE subfamily. The asterisk indicates the presence of the corresponding TF motif in the TE consensus sequence.

| Species | Genome version | # MER81 |
|---|---|---|
| human | hg19 | 3668 |
| marmoset | calJac3 | 3148 |
| rhesus monkey | rheMac3 | 3497 |
| bushbaby | otoGar3 | 2604 |
| tarsier | tarSyr2 | 3357 |
| tree shrew | tupBel1 | 1628 |
| lemur | micMur1 | 2759 |
| mouse | mm10 | 505 |
| rat | rn5 | 498 |
| prairie vole | micOch1 | 516 |
| rabbit | oryCun2 | 1581 |
| bat | myoLuc2 | 2213 |
| megabat | pteVam1 | 2710 |
| hedgehog | eriEur2 | 534 |
| dog | canFam3 | 2891 |
| cat | felCat5 | 3124 |
| cow | bosTau7 | 2758 |
| pig | susScr3 | 2842 |
| alpaca | vicPac2 | 3116 |
| horse | equCab2 | 3895 |
| elephant | loxAfr3 | 3292 |
| armadillo | dasNov3 | 3156 |

| | | |
|---|---|---|
| dolphin | turTru2 | 3403 |
| opossum | monDom5 | 0 |

**Supplemental Table 1.** MER81 annotations by RepeatMasker in 24 selected mammals. These mammals cover major modern lineages. The numbers of MER81 copies in each species (each specific genome build) were obtained from RepeatMasker annotations.

| Target | Distance to target TSS | Target gene annotation |
|---|---|---|
| *IFNAR1* | 58 kb | Surface receptor of IFN-α signaling |
| *TYK2* | -5 kb | Downstream kinase required for IFN-α signaling transduction |
| *IFNGR2* | -19 kb | Surface receptor of IFN-γ signaling |
| *IRF1* | 15 kb | Known transcription factor activated by IFN-γ signaling |
| *JAK2* | -47 kb | Downstream kinase required for IFN-γ signaling transduction |
| *CAMK2B* | -131 kb | Phosphorylation for IFN signaling member STAT1 in response to IFN-γ |
| *RAP1B* | -130 kb | A membrane protein inducible by IFN-γ signaling |
| *CX3CL1* | 9 kb | Chemokine to recruit leukocytes |
| *CXCR5* | -0.3 kb | Involved in B-cell maturation and migration |

**Supplemental Table 2.** Examples of RELA bound MER81 instances near interferon signaling-related genes. These instances were identified with RELA binding from HAEC RELA ChIP-seq data. Highlighted were the distance from instances to gene TSS and notes for the gene relevance to interferon signaling.

**qPCR Primers**

| | |
|---|---|
| hIFNAR1_primary_1F | TCCCTGGAGCTGGACTAGTC |
| hIFNAR1_primary_1R | TGTGAGGCCAAGTGTAGCAG |
| Tub1a_F | GTCCAACACGAGGTCAATGAT |
| Tub1a_R | GCAACTTATCACAGGCAAAGAAG |
| IFNGR2_intronic_F | CCTGGCAGAATCCAGAAGAG |
| IFNGR2_intronic_R | GTGCCCTCGAGTGAAGAAAG |

**gRNAs**

| | | |
|---|---|---|
| Mer81 | gRNA_L | CCGGGACAGTACTAGATTAG |
| Mer81 | gRNA_R | TAACCCGTCTACAAGCATCA |
| subMer81 | gRNA_L | ACTTATCCAGAGCACGAGGCAGG |
| subMer81 | gRNA_R | CTGGTTTGTCAGCGACTCAGGGG |

**Supplemental Table 3.** Primer sequences and gRNAs.
